# DNA methylation enables recurrent endogenization of giant viruses in an animal relative

**DOI:** 10.1101/2024.01.08.574619

**Authors:** Luke A. Sarre, Iana V. Kim, Vladimir Ovchinnikov, Marine Olivetta, Hiroshi Suga, Omaya Dudin, Arnau Sebé-Pedrós, Alex de Mendoza

## Abstract

5-methylcytosine (5mC) is a widespread silencing mechanism that controls genomic parasites. However, in many eukaryotes 5mC has gained complex roles in gene regulation beyond parasite control. Animals are a quintessential case for 5mC evolution, as they show widespread variability across lineages, ranging from gene regulation and transposable element control to loss of this base modification. Here we show that the protist closely related to animals *Amoebidium appalachense* features both transposon and gene body methylation, a pattern reminiscent of invertebrates and plants. Unexpectedly, large hypermethylated regions of the *Amoebidium* genome derive from viral insertions, including hundreds of endogenized giant viruses contributing 14% of the encoded genes, to an extent never reported before in any eukaryotic genome. Using a combination of inhibitors and functional genomic assays, we demonstrate that 5mC silences these giant virus insertions. Moreover, alternative *Amoebidium* isolates show polymorphic giant virus insertions, highlighting a dynamic process of infection, endogenization and purging. Our results indicate that 5mC is critical for the controlled co-existence of newly acquired viral DNA into eukaryotic genomes, making *Amoebidium* a unique model to understand the hybrid origins of eukaryotic genomes.

## Main Text

5-methylcytosine (5mC) is a common base modification among eukaryotes(1–3). 5mC is deposited by DNA methyltransferases, a family of enzymes with ancestral families conserved throughout eukaryotes(4, 5). Some DNMTs are maintenance type enzymes, perpetuating 5mC patterns, including DNMT1 and DNMT5, while other DNMTs have *de novo* activity, such as DNMT3(6, 7). However, the DNMT repertoire of an organism is not predictive of 5mC function. In some eukaryotes, including plants and animals, 5mC is associated with gene regulation, exemplified by gene body methylation, where 5mC positively correlates with gene transcriptional levels(1, 3, 8). However, the most widespread role of 5mC is in transposable element (TE) silencing, which is the assumed ancestral role in eukaryotes(9, 10).

Despite most attention being devoted to controlling endogenous parasitic elements, one of the first described functions of 5mC in eukaryotes was to silence retroviral insertions in mammals(11). Similarly, in bacteria the main role of 5mC is to combat viruses(12). Therefore, controlling exogenous viral invasions is arguably as important as TE control for epigenetic silencing. Interestingly, it is increasingly recognised that many eukaryotic genes have viral origins, co-opted repeatedly throughout evolution(13). One of the most common sources for these acquisitions are giant viruses (Nucleocytoviricota). Giant viruses have a wide range of eukaryotic hosts and are present in almost all ecosystems, posing a widespread threat to eukaryotic cells(14, 15). Giant viruses are exceptional among viruses as they possess enormous genomes (100 kb to 2.5 Mb) encoding many proteins thought to be eukaryotic hallmarks such as histones(14, 15). Giant viruses originated before modern eukaryotes, and they are believed to have contributed essential genes to eukaryogenesis(16). Furthermore, recent reports indicate that giant viruses can endogenize into extant eukaryotes(17–19). However, how this potentially lethal DNA is incorporated into eukaryotic genomes is currently not understood.

Finding a link between viral control and epigenetic regulation however, is hampered by the scarcity of reported recent giant virus endogenizations(17, 20). Moreover, 5mC is evolutionarily very plastic, and many eukaryotic lineages have lost this epigenetic modification(1, 2), possibly because of its mutagenic potential and cytotoxic off-target effects of DNMTs(21). Furthermore, 5mC function varies across lineages. In fungi 5mC is restricted to silencing TEs(22), whereas in invertebrates 5mC is usually restricted to gene bodies, and most TEs remain unmethylated(1, 2, 23). To expand our knowledge of 5mC systems and to unravel how a potentially ancestral fungal-like methylation pattern gave rise to the animal 5mC system, we focused on protists of the holozoan clade. These close animal relatives form four major lineages: choanoflagellates, filastereans, ichthyosporeans and pluriformeans (**Fig. 1A**)(24, 25). In recent years, unicellular holozoan genomes have been shown to encode many genes previously thought to be unique to animals, informing the complex genomic nature of the unicellular ancestors of animals(24–27). However none of these genomes encode DNMTs, suggesting an evolutionary loss of 5mC capacity(28). In this study, we fill this gap by describing a unicellular relative of animals that has maintained 5mC, and unexpectedly find an unappreciated and potentially ancestral use of 5mC in regulating giant virus endogenizations.

## Results

### The *Amoebidium* genome presents both gene body and TE methylation

To reconstruct the pre-animal roots of 5mC, we searched the available genomes and transcriptomes of unicellular holozoans for DNMT1 orthologues(27, 29), the maintenance DNMT in animals. We discovered that DNMT1 is expressed by *Amoebidium appalachense* (**Fig. 1A**), an ichthysoporean originally isolated from the cuticle of freshwater arthropods that can be grown axenically(30). Like other Ichthyosporea(31), *Amoebidium* are parasites or commensals of animal hosts, and show a coenocytic life cycle, with cells dividing their nuclei in a shared cytoplasm until forming a mature colony, that gives rise to unicellular and uninucleated cells(32). Yet, *Amoebidium* branches quite deep within the ichthyosporean lineage (**Fig. 1A**). Interestingly, *Amoebidium* DNMT1 sequence presents the same domain architecture of animal DNMT1 orthologues, including a zinc finger CXXC absent in non-holozoan sequences (**fig. S1A**), which suggests this domain architecture was an innovation of holozoans yet DNMT1 was lost repeatedly.

**Fig. 1.**
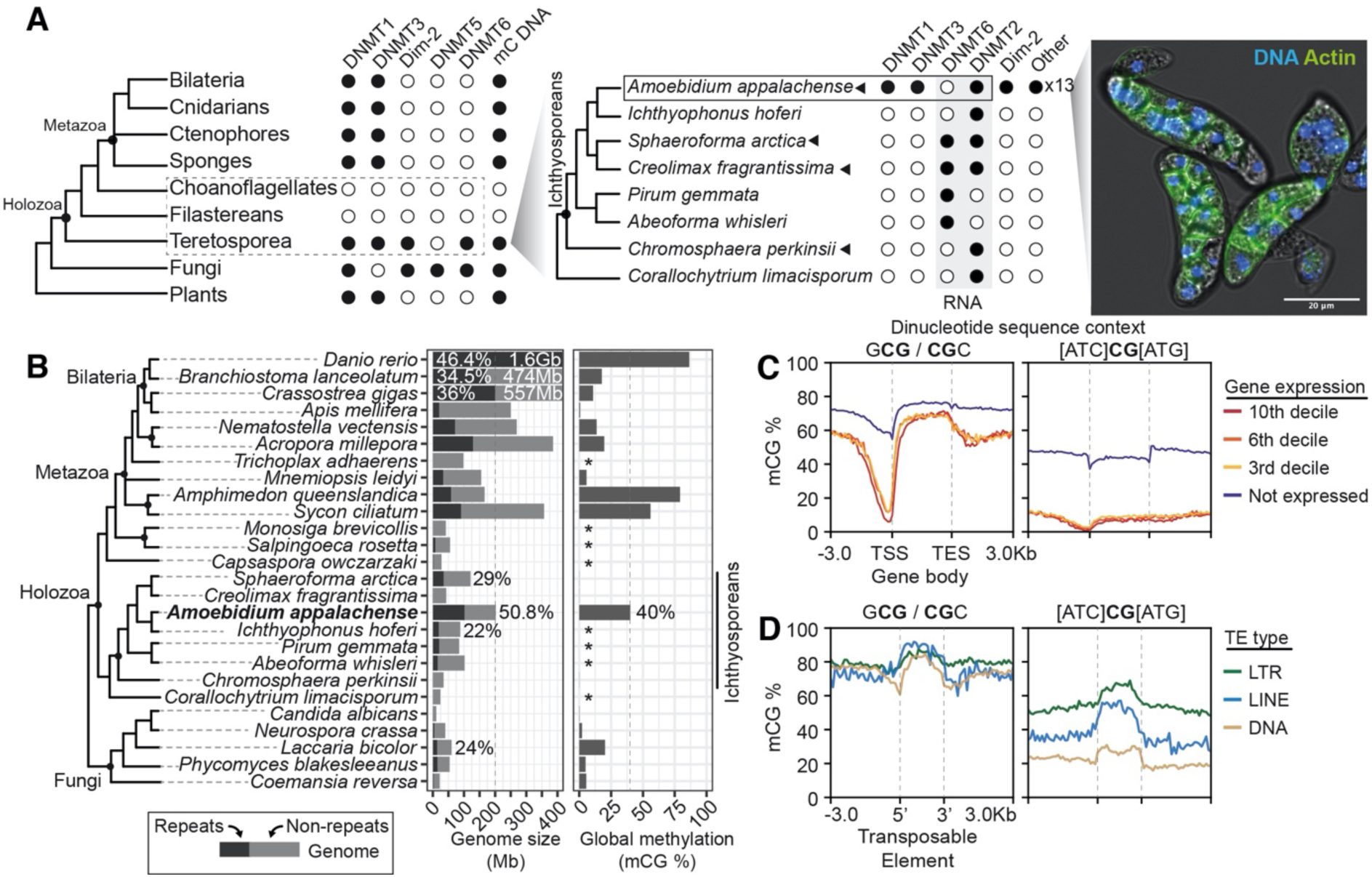
*Amoebidium appalachense* displays a methylome with gene body and transposon methylation. (**A**) Distribution of DNA Methyltransferases in eukaryotes and teretosporeans. Black dot indicates presence, and white dot indicates absence, black triangles species for which Enzymatic Methyl-seq has been performed. *Amoebidium* cells are stained with phalloidin and Hoescht. Phylogenetic relationships based on previous studies(27). (**B**) Distribution of genome sizes, repetitive content and global methylation levels across opisthokonts. Asterisks indicate species for which the lack of 5mC is inferred from their absence of DNMTs. Genomes sizes for species with genomes above 400Mb are displayed in white, and repeat content (%) is also highlighted for clarity. (**C**) Gene body methylation and (**D**) Transposable element (TE) methylation averages split by the extended CG sequence context. Not expressed genes display <1 TPM. TSS: Transcriptional Start Site; TES: Transcriptional End Site. TEs only include copies spanning at least 70% of the RepeatModeler2 consensus model.

To fully characterise the gene repertoire and investigate *Amoebidium* 5mC patterns, we sequenced the genome of this species using a combination of nanopore long reads, Illumina short-reads and micro-C. We obtained a chromosome-scale assembly spanning 202 Mb, with 96.6% of the sequence contained in 18 chromosomes and a 95% BUSCO score (**Fig. 1b**, **fig. S2, A and B**). When examining *Amoebidium*’s DNMT repertoire, we identified a total of 18 DNMTs, comprising orthologues of DNMT1, DNMT3, DNMT2, Dim-2, and various lineage-specific clades (**Fig. 1A**, f**ig. S1, B and C**). Notably, unlike DNMT1, none of the other DNMTs exhibit additional domains. Specifically, DNMT3 lacks the protein domains found in animal DNMT3, such as PWWP or ADD (**Fig. S1A**)(7, 28). In contrast, none of the ichthyosporeans with genomic data available show DNMTs other than the RNA-specific DNMT2 and DNMT6 (**Fig. 1A**)(3, 7, 33). Thus, *Amoebidium* is the only sequenced unicellular holozoan species that retains the ancestral eukaryotic complement of DNMTs, highlighting the pervasive tendency of eukaryotes to lose 5mC.

Next, we performed whole-genome DNA methylation profiling to analyse the 5mC patterns in *Amoebidium*, as well as in three other ichthyosporean species lacking DNA DNMTs as negative controls. In *Amoebidium*, global methylation levels soar to 40%, exclusively within the CG dinucleotide context, setting it apart from most invertebrates and fungi and the other ichthyosporean species, which exhibit negligible levels of 5mC(**Fig. 1B**)(1, 22). Notably, not all CG dinucleotides exhibit uniform methylation levels. Specifically, the symmetrical m<u>CG</u>C and Gm<u>CG</u> trinucleotides stand out with hypermethylation levels at around 70%, whereas the remaining CG dinucleotides maintain lower levels at approximately 20% (f**ig. S3A**). This suggests that *Amoebidium* boasts elevated methylation levels with a wider sequence specificity beyond the CG dinucleotide, a context-dependent regionalisation of 5mC reminiscent of heterochromatin methylation in mammals(34), likely reflecting the sequence preferences of the diverse *Amoebidium* DNMTs.

Considering the high global methylation levels in *Amoebidium*, we proceeded to investigate which genomic regions exhibit enriched 5mC. Protein coding genes displayed a gene body methylation pattern reminiscent of plants and animals, with relatively low levels of promoter methylation (**Fig. 1c**)(35, 36). However, *Amoebidium*’s gene body methylation is not positively correlated with transcription as in plants or animals, as all active genes have similar methylation levels irrespectively of transcriptional level, whereas silent genes show higher methylation, including the promoter (**Fig. 1c, fig. S3B**). Therefore, gene body methylation appears to predate animal origins in the holozoan clade, yet its positive association with transcription was an animal innovation potentially linked to the domain acquisitions of animal DNMT3s (f**ig. S1A**).

In contrast to many invertebrates(1, 2), *Amoebidium* exhibits targeted methylation of TEs (**Fig. 1D**, **fig. S3, B and C**). Notably, methylation levels are highest in recent TE insertions and on transcriptionally silent genes (**Fig. 1D, fig. S3C**), regardless of the adjacent CG sequence context. In contrast, gene body methylation of actively transcribed genes primarily occurs within the CGC/GCG trinucleotide context (**Fig. 1C**). This indicates that in *Amoebidium*, the 5mC link with gene silencing is exclusive to methylation of all CG contexts beyond the widespread methylation of the CGC/GCG trinucleotide. Further supporting the link between 5mC and TE silencing, approximately 50% of *Amoebidium*’s genome is comprised of TEs, a level unmatched in any unicellular holozoans, yet similar to vertebrates such as humans (50%) or zebrafish (**Fig. 1B, fig. S2, C and D**). Therefore, the genome of *Amoebidium* is permissive to TE expansions because 5mC can silence these novel insertions by reducing their potential deleterious effects, similar to what has been proposed for vertebrates(37),

### Large hypermethylated regions uncover multiple giant repeats including hundreds of giant virus insertions

To characterise the chromosome-level distribution of 5mC, we took advantage of the relative depletion of non-CGC/GCG methylation to locate regions of hypermethylation across the genome. We found many islands of hypermethylation spread across the chromosomes (**Fig. 2A**), many of which were consistent with regions of high TE content. However, many presented highly gene-rich areas spanning up to 200 kilobases, with most genes showing few to no introns, in clear contrast to the exon-rich *Amoebidium* genes (average 7.2 exon/gene, **fig. S4A**). Further characterization of these areas revealed core giant virus genes, including Poxvirus Late Transcription Factor (VLTF3), A32-like packaging ATPase, D5 DNA primase, or NLCDV major capsid proteins (**fig. S4A**)(14, 38). Using these core genes, we searched the NCBI database and performed phylogenetic analyses using curated databases of giant viruses marker genes(38). We found that these insertions could be classified as belonging to a lineage member of the Pandoravirales, closely related to Medusavirus or Clandestinovirus (infecting amoebozoans, **fig. S5, A and B**). Yet not all pandoraviruses in the *Amoebidium* genome originate from a single catastrophic insertion event or even a single viral lineage, as they show high levels of sequence divergence among them (**fig. S5C**). Furthermore, there are insertions in almost all chromosomes (90 chromosomal insertions, with 42 sequences in unplaced contigs), some having accumulated secondary TE insertions (**Fig. 2A**). The disparity of insertion lengths, and the observation that none of them encode a full repertoire of core giant virus genes (**Fig. 2B**), suggests that complete viral genome integrations are rare, or that gene loss occurs rapidly after insertion.

**Fig. 2.**
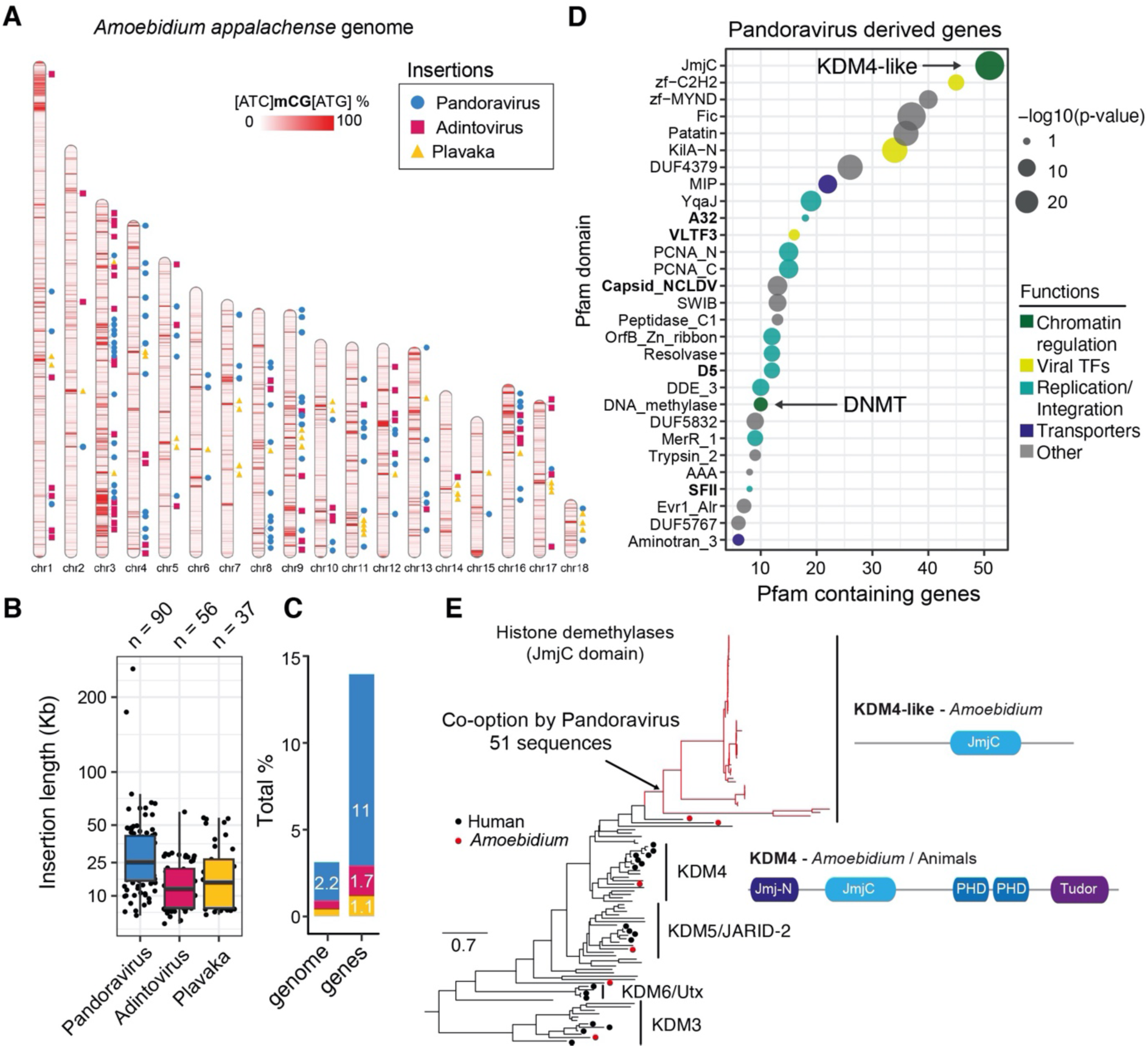
The *Amoebidium* genome harbours hundreds of viral insertions. (**A**) Location of giant virus (Pandoravirus), adintovirus and Plavaka giant repeats across the *Amoebidium* genome. Windows of 10kb are coloured according to their methylation level in non-CGC/GCG trinucleotides. (**B**) Distribution of insertion sizes of the giant repeats within chromosomes. Centre lines in boxplots are the median, box is the interquartile range (IQR), and whiskers are the first or third quartile ± 1.5× IQR. (**C**) Contribution of giant repeats to genome size and gene counts. (**D**) Pfam domains enriched in genes encoded in pandoravirus endogenized regions. In bold, marker giant virus domains. **e**) Maximum-likelihood phylogeny of JmjC in eukaryotes, highlighting the expansion of KDM4-like enzymes in pandoravirus-derived regions. Black dots indicate human sequences, red dots indicate *Amoebidium* sequences, red branches indicate genes within endogenized viral regions. Domain architectures defined with PFAM domains.

In addition to the pandoraviruses, other compact hypermethylated regions were characterised by genes encoding VLTF3, Dam methyltransferase, minor and major capsid proteins, and a DNA polymerase family B (**fig. S4B**). DNA polymerase sequences produced closest matches to adintovirus, a group of recently described double stranded DNA Polinton-related viruses thought to exclusively infect animals(39)(**fig. S5D**). It is worth noting many Polinton-related viruses are virophages or descendants of these(40, 41), known to parasitise giant viruses, which could explain the abundance of these sequences in the *Amoebidium* genome. Similarly to the pandoraviruses, not all adintoviruses were closely related among each other (**fig. S5E**), suggesting multiple independent insertion events. In contrast to pandoraviruses, some insertions kept long terminal repeats and were complete (∼30kb, **Fig. 2B**), yet others were truncated and in the process of degeneration.

Then, we identified a third type of giant repeat, consisting of tandem clusters of repetitive intron-poor genes up to 50 kb long, usually flanked by a Plavaka transposase (**fig. S4C**)(42). Many of these genes encode for tyrosine recombinases, and interestingly their only hits in the NCBI NR database belong to very distant eukaryotic lineages including dinoflagellates or red algae, thus suggesting some form of lateral gene transfer as their source (**fig. S4C**). When we combine the three types of highly methylated giant repeats, they make up 3.1% of *Amoebidium*’s total DNA. Surprisingly, their contribution to the protein-coding genes constitutes 14% of the entire proteome, with the majority originating from viruses. The amount of giant virus insertions in *Amoebidium* is the largest ever reported in any eukaryote (**Fig. 2C**).

### Endogenized giant virus co-opted eukaryotic histone-demethylases

To understand the potential contribution of endogenized genes to the *Amoebidium* gene repertoire, and also to better understand the gene complement of the original pandoravirus genomes, we characterised the functional enrichment of genes encoded in these endogenized regions. An enrichment test of Pfam domains revealed many domain categories involved in the viral replication and integration process (recombinases, integrases, PCNA), viral gene regulation (Transcription Factors), or some transporters (e.g. Aquaporins/MIP), which are likely critical to taking control of the host during infection (**Fig. 2D**)(43). Gene ontologies also suggested these genes were enriched in membrane fission or tubulin depolymerization (**fig. S6A**). Notably, some of the most enriched categories were involved in chromatin regulation. Among these 10 out of the 18 DNMTs encoded in the *Amoebidium* genome reside in pandoravirus insertions, which suggests that these could be used by the virus to modify its own DNA. Consistently, giant viruses, and pandoravirus in particular, are known to use various forms of DNA methylation (N6-methyl-adenine and N4-methyl-cytosines) to methylate their own genomes(44), which might play a role in infection. However, the *Amoebidium* pandoravirus DNMTs form a sister group to other giant virus uncharacterised DNMTs (**fig. S1B**), thus they were not recently acquired from the host and their sequence-substrate preferences remain unknown.

The most enriched endogenized domain is the Jumonji C (JmjC) domain. Although JmjC domains can perform many enzymatic functions, our phylogenetic analysis revealed that these are divergent paralogues of the Histone lysine demethylase subfamily 4 (KDM4). Notably, despite JmjC containing proteins have been identified in giant viruses(45), we could not find any KDM4-like JmjC homologues in publicly available giant virus genomes. *Amoebidium* encodes a canonical KDM4 orthologue like those of other eukaryotes, including its characteristic histone interacting domains (PHD, Tudor, **Fig. 2E**). However, the endogenized KDM4-like enzymes only contain the enzymatic JmjC domain (**Fig. 2E**). KDM4 enzymes are known to demethylate histone 3 tail lysines, most commonly lysine 9 (H3K9me2/3) or lysine 36 (H3K36me2/3) residues. Although many giant viruses encode all four eukaryotic nucleosome histones (H2A/B,H3,H4)(38, 46), we did not find any in the viral insertions. Furthermore, viral histones present very divergent histone tails(47), thus it is unlikely that KDM4-likes are used to control potential pandoravirus histones. Instead, given the conserved role of H3K9me3 in heterochromatin formation in eukaryotes, KDM4-like enzymes could be used by the virus to avoid silencing by the host chromatin. In KDM4 overexpressing cancer cells, depletion of H3K9me3 promotes DNA breaks and genome instability(48), a process which could serve the virus to integrate into the host genome, or explain the amount of endogenization events.

The KDM4-like enzymes stand out among the endogenized genes as they have preserved the multi-exon domain structure of eukaryotic genes (**fig. S6C**), unlike the vast majority of giant virus genes that lack introns. While most of the endogenized genes remain silent in culture conditions, four of these KDM4-like genes are transcribed (TPM >1)(**fig. S6C**). Moreover, JmjC genes are found in 39% (52) of the insertions, which could reflect a lower chance of purging those genes after the insertion event. Intriguingly, a couple of KDM4-like genes are found outside hypermethylated giant virus regions and are flanked by normal host genes, showing almost exclusive mCGC/GmCG methylation (the default state for transcribed host genes, **fig. S6C**). Given their basal position in the phylogeny of KDM4-likes (**Fig. 2E**), these genes could be *Amoebidium*-specific KDM4 divergent paralogues that already lost the chromatin interaction domains compared to the canonical KDM4 copy, and were later co-opted by the pandoraviruses. Alternatively, giant virus might have originally acquired a canonical KDM4 gene from the host, which then lost some of its companion domains to perform virus associated functions. Then, these basally branching KDM4-like genes are the remnants of past pandoravirus insertions, where most other viral genes have been purged and only JmjC loci are kept, being domesticated to become part of the host repertoire. Thus, the intricate interaction between the host chromatin and the giant viruses is likely critical to explain the gene flow between the host and parasite.

### DNA methylation removal is sufficient for viral transcriptional reactivation

Since dense 5mC demarcates the viral insertions and these are transcriptionally silent, we wanted to directly investigate the causal relationship between 5mC and gene expression in *Amoebidium*. We tested the effect of cytidine-analogues 5-Azacytidine, Zebularine and Decitabine (which block DNMTs and lead to passive dilution of 5mC(49, 50)) to investigate the impact of 5mC on gene expression. A 3-day cytidine-analogue treatment spans at least two generations of *Amoebidium* colonies, covering two rounds of coenocytic development starting from an uninucleate cell to colony maturation and cell release in ∼30 hours (**fig. S7A**). Therefore, several rounds of nuclear division maximises the potential of obtaining sufficient passive 5mC loss. Importantly, 5mCG remains constant across development, thus minimising the potential confounding staging effects across treatments (**fig. S7, B and C**). We then used Enzymatic Methyl-seq to quantify 5mC of the treated cells and found that only 5-Azacytidine showed a decrease in global methylation levels (from ∼40 to 15%, **Fig. 3A**). Consistently, only the *Amoebidium* cells treated with 5-Azacytidine showed growth defects and increased mortality (**fig. S7D**). However, 5-Azacytidine can potentially be incorporated into RNA and be cytotoxic(51, 52). To control for those off-target effects, we also treated two ichthyosporean species lacking genomic 5mC with 5-Azacytidine, observing mild growth defects (**fig. S7E**).

**Fig. 3.**
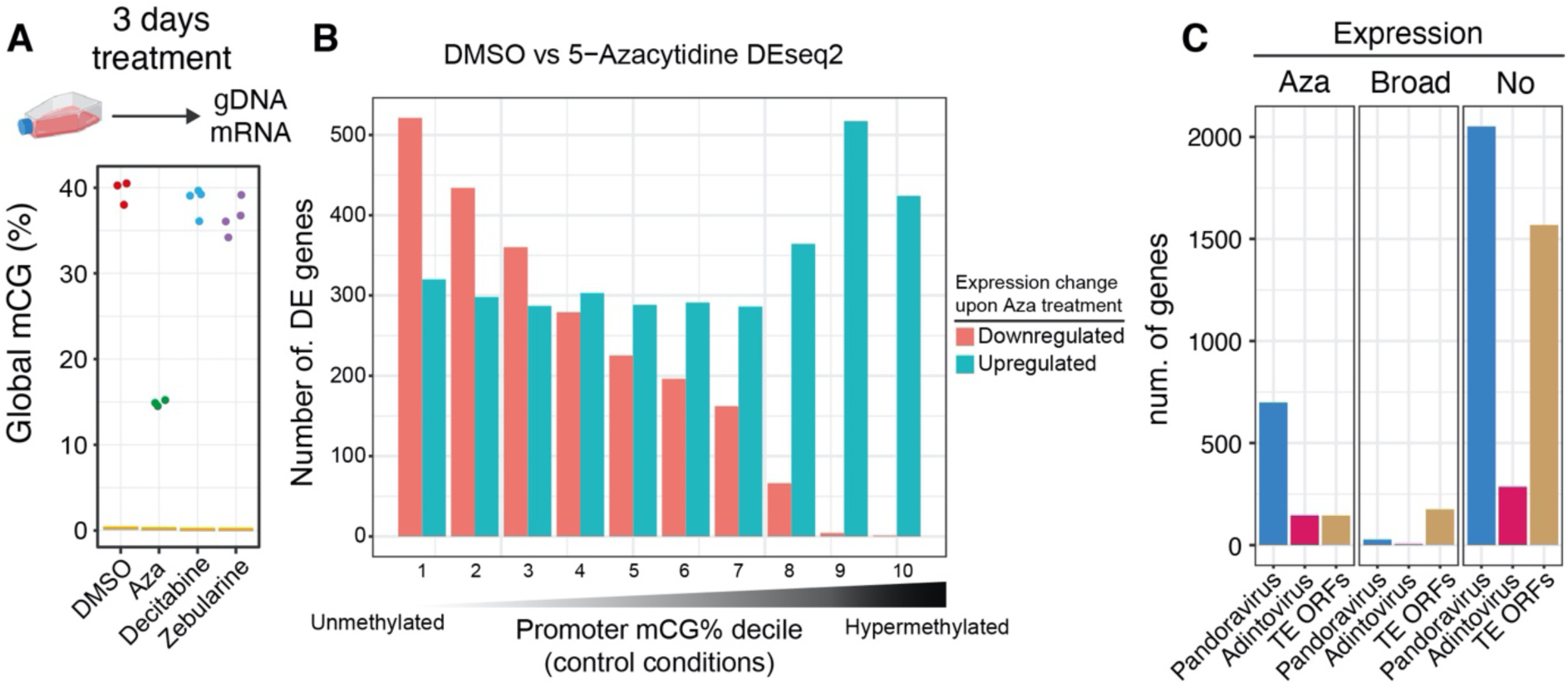
5mC removal leads to viral transcriptional reactivation. (**A**) Global methylation levels measured with Enzymatic Methyl-seq for *Amoebidium* cultures treated for 3 days with DMSO, 5-Azacytidine (Aza), Decitabine and Zebularine. (**B**) Distribution of differentially expressed genes classified according to the promoter methylation status in untreated conditions (divided in deciles). The bar colour depicts the direction of change upon 5-Azacytidine treatment. (**C**) Number of genes encoded in pandoraviruses, adintoviruses or TE Open Reading Frames according to their transcriptional response to 5-Azacytidine treatment. “Aza” are genes that are only transcribed upon treatment, “Broad” are genes that are expressed in any moment of *Amoebidium* development / control conditions, and “No” are genes that are not expressed in any condition (TPM <1).

We then used RNA-seq to characterise the transcriptional response to 5-Azacytidine in *Amoebidium* and *Sphaeroforma arctica*. *Sphaeroforma* is a closely related ichthyosporean that also possesses a relatively large amount of TEs and few instances of polinton-type viruses(53), yet lacks genomic 5mC (**Fig. 1B**). Both species showed hundreds of differentially expressed genes upon treatment (5630 in *Amoebidium* and 1807 in *Sphaeroforma*, fdr < 0.01), but very few of these showed consistent dynamics across species (**fig. S8A**), thus not suggesting generic stress response shared across species. Nevertheless, genes that upregulated upon 5-Azacytidine in *Amoebidium* have stress associated Gene Ontologies, while a wide range of metabolic processes are downregulated (**fig. S8D**). As observed in 5-Azacytidine treated cancer cells, the stress response might be driven by TE reactivation(54). Focusing on TEs, only *Amoebidium* showed a drastic expression increase in almost all TE types after 5-Azacytidine (**fig. S8B**), whereas *Sphaeroforma* did not show any particular enrichment in TE or viral upregulation (most remaining transcriptionally silent / unchanged, **fig. S8B**), suggesting that the TE response to 5-Azacytidine is a direct consequence of 5mC loss. We further validated this observation by dividing *Amoebidium* genes according to their promoter methylation level in untreated conditions. Genes that normally present unmethylated promoters had a mixed transcriptional response to DNA methylation removal suggestive of indirect effects, whereas genes with hypermethylated promoters were almost exclusively upregulated upon 5-Azacytidine treatment (**Fig. 3B**). Thus, 5mC is a silencing mark in *Amoebidium* sufficient to repress methylated genes.

When inspecting Giant Virus and Adintovirus endogenized genes, we saw a consistent transcriptional reactivation upon methylation removal. 737 genes encoded in pandoravirus insertions (26%) were transcriptionally reactivated (**Fig. 3C**), with the majority of them being the JmjC genes, but also many genes involved in gene regulation (**fig. S8C**). Similarly, 144 adintovirus genes were reactivated upon demethylation (32%). However, we did not observe formation of viral particles through microscopy, and consistently we did not see transcriptional reactivation of capsid proteins. This suggests that viral formation would require extra genes that might have been purged or have accumulated critical mutations since the insertion occurred. Alternatively, post-transcriptional silencing mechanisms could stop the formation of mature viral particles. In sum, direct manipulation of the host methylome demonstrates that 5mC is instrumental for silencing and minimising the consequence of viral DNA acquisition.

### Giant virus endogenization is polymorphic and highly dynamic in *Amoebidium*

Maintaining a substantial quantity of potentially harmful viral DNA in the *Amoebidium* genome may serve as an adaptive mechanism with significant roles. Conversely, it could also represent a passive outcome facilitated by epigenetic silencing. To assess these hypotheses, we set out to compare genetically distinct *Amoebidium* isolates from our reference genome. We first obtained the transcriptome of 6 isolates, 4 belonging to *A. appalachense* and 2 to *A. parasiticum*. Whereas the isolate’s 18S sequences were identical at the species level (**fig. S9A**), the rapidly divergent mitochondrial 16S revealed 4 clades, including a slightly divergent *A. appalachense* lineage (**Fig. 4A**). We selected a member of the divergent *A. appalachense* lineage (Isolate 9181) and one *A. parasiticum* (Isolate 9257) for genome sequencing using nanopore long reads. Genome assembly size varied across isolates (**Fig. 4A**), yet annotation qualities and nanopore-assessed 5mC levels were consistent (**fig. S9B**).

**Fig. 4.**
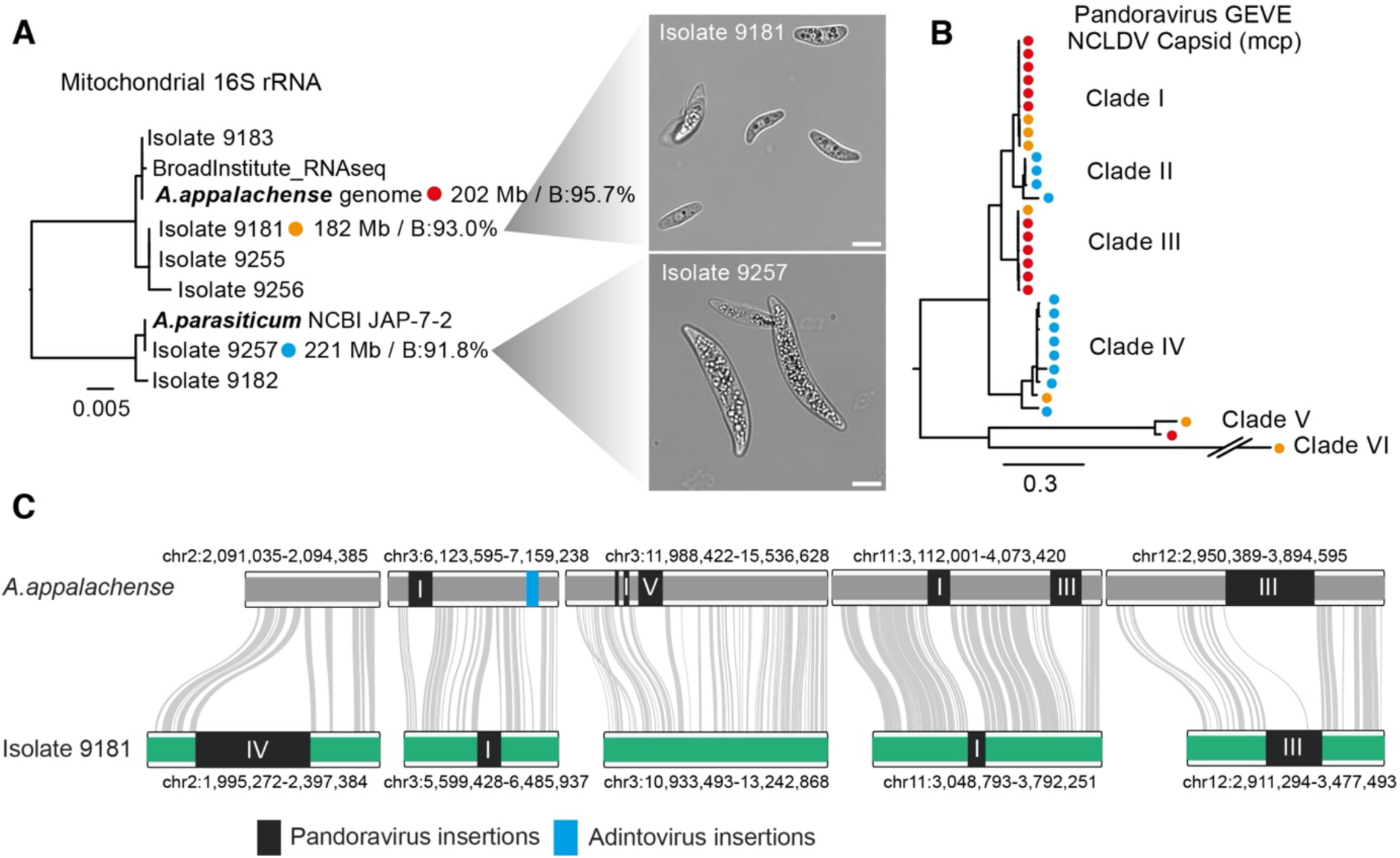
*Amoebidium* isolates display rapid turnover of viral endogenization events. (**A**) Maximum likelihood phylogenetic tree displaying the mitochondrial 16S phylogeny of *Amoebidium* isolates. The reference genome (red), Isolate 9181 (orange) and Isolate 9257 (blue) display the genome assembly characteristics: size (megabases) and BUSCO completeness (B %). Micrographs display *Amoebidium* isolate cells in culture, with the white bar spanning 10 µm. (**B**) Maximum likelihood phylogeny of NCLDV Major capsid proteins encoded in pandoravirus endogenization events. Dots represent the genome they come from following panel **A** colour code. Double slash indicates that the branch has been shortened for display purposes. (**C**) Conserved and polymorphic viral insertions across reference genome and Isolate 9181. Grey lines indicate presence of one-to-one orthologues, whereas dark rectangles indicate viral insertions. Roman numerals indicate the clade of the pandoravirus according to panel **B** phylogeny.

Annotation of the viral endogenizations in these alternative genotypes revealed a dynamic and diverse history for pandoraviruses and adintoviruses associated with the *Amoebidium* lineage. Phylogenetic markers such as the major capsid proteins or VLTF3 revealed that at least 6 separate clades of pandoraviruses infect these protists, with some clades unique to one isolate (Clade II) and others shared by the isolates (Clade IV) but absent in the reference genome (**Fig. 4B, fig. S9C**). Similarly, four adintovirus clades are found across the isolates, with some being shared across all 3 genomes (**fig. S9D**). Notably, Isolate 9257 shows only 4 adintoviruses compared to the 44 present in the reference genome. This reveals that viral diversity infecting *Amoebidium* is not limited to a single lineage and is often endogenized in an isolate-specific manner.

Since sequence similarity and gene synteny between Isolate 9181 and the reference genome remains highly conserved (**fig. S9E**), we used this to assess the ancestral nature of endogenization events. Despite their close phylogenetic relationship, only a minority of endogenization events were shared across the isolates, with most featuring polymorphic insertions amidst synteny blocks (**Fig. 4C**). Giant viruses are not known to require integration into the host genome during their infectious cycles(14), this process appears to be stochastic, potentially occurring during unsuccessful infections and at various chromosome positions. Additionally, it underscores the dynamic nature of integration, which the host tolerates through multiple cycles, with most endogenized elements being quickly eliminated after insertion.

## Discussion

Here we show how a unicellular eukaryote closely related to animals undergoes a recurrent process of mixing its genome with that of its giant virus predators. This foreign DNA is curbed by 5mC silencing, allowing for survival after these potentially lethal events. We propose that epigenetic silencing greatly reduces the lethality of these endogenization events. Supporting this general hypothesis, many of the previously described large-scale giant virus endogenizations in eukaryotes, including early land plant lineages, green algae or the amoebozoan *Acanthamoeba castellanii*, coincide with species that have retained 5mC as a silencing mechanism (**Fig. 5**)(17–19, 55, 56). Plants that are not dependent on water for reproduction (Spermatophyta) or germline-segregating animals are likely protected from giant virus endogenization events despite carrying silencing mechanisms(18), yet chromosome-scale genomes and directed searches might reveal exceptions to this rule. Giant viruses infect all kinds of eukaryotic groups, but eukaryotes that have secondarily lost 5mC silencing, as exemplified in all the other available unicellular holozoan genomes, rarely present large giant viral DNA insertions. Notably, most eukaryotic clades have genes derived from giant viruses(13), or insertions of double-stranded medium size DNA viruses like polinton-like/virophages(53), suggesting that infection and endogenization are widespread, but the retention of these insertions is uneven across lineages. It is likely that other silencing mechanisms other than 5mC, such as histone modifications or small RNAs, can be used for this purpose(57–59), yet arguably the cost of integrating large amounts of viral DNA in epigenetically unprotected species is probably coped with by extremely rapid purging or take over by uninfected conspecific cells.

The 5mC patterns in *Amoebidium* also suggest that gene body methylation predates animal origins. Although this would potentially support the hypothesis that gene body methylation was present in the ancestor of eukaryotes(35), this pattern is very sparsely distributed (**Fig. 5**). Among chlorophytes, only *Chlorella variabilis* has a pattern similar to that of *Amoebidium*(35), while absent in *Chlamydomonas* and Prasinophytes(5, 36). Spermatophytes (Angiosperms(60), conifers(61) and ferns(62)) show gene body methylation, whereas liverworts and mosses generally lack it(35, 63, 64). In contrast, the more basally branching streptophyte *Klebsormidium nitens* shows gene body methylation(55), albeit in a pattern quite divergent to land plants, which could suggest independent origins of gene body methylation in these lineages. Interestingly, *Amoebidium*, *Chlorella* (65) and *Klebsormidium* (55) present giant virus endogenization events, which suggests that gene body methylation might arise as a convergent response or by-product to recurrent infections and expansion of parasitic DNA(60), perhaps avoiding intra-genic elements hijacking transcription from the host genes(66). In particular, *Amoebidium* encode DNMT3 and an animal-like DNMT1, which could support that gene body methylation across holozoans is homologous and deposited by orthologous DNMTs. The link with gene body and transcription became an animal innovation through the acquisition of PWWP and ADD domains in animal DNMT3 orthologues, starting a feedback loop with histone modifications such as H3K36me2/3(67–69). Regulation of host gene transcription in multicellular animals might have restricted and weakened the role of 5mC in TE silencing, suggested by its absence across many invertebrate genomes(1, 2).

**Fig. 5.**
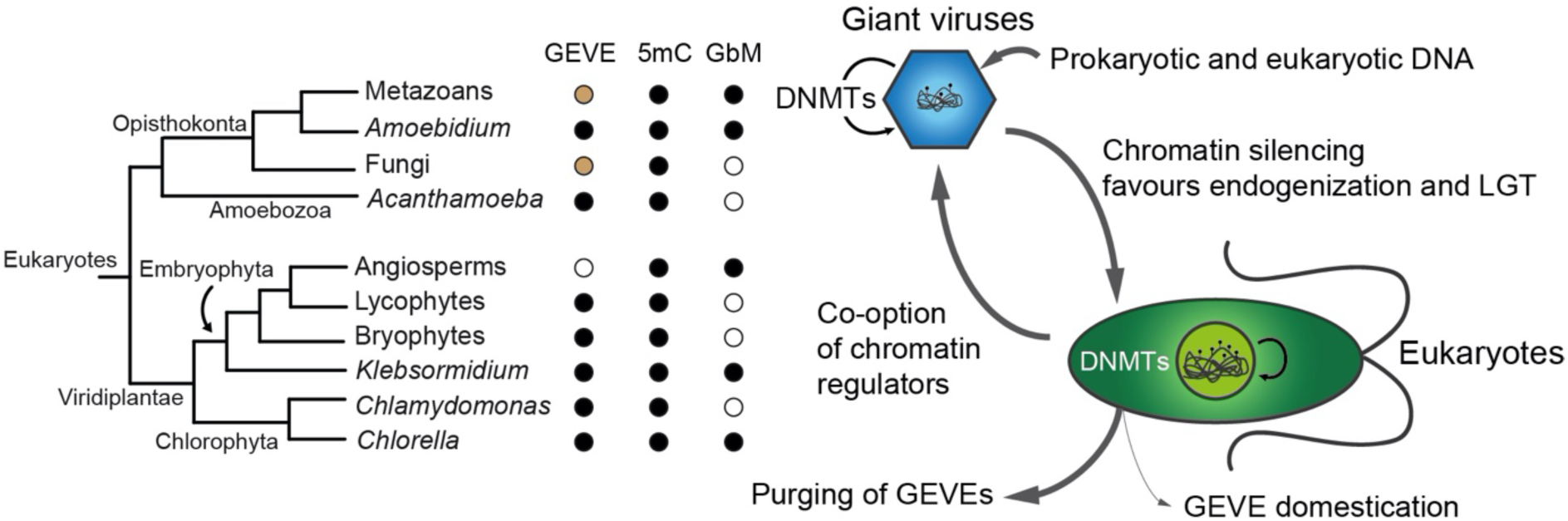
Giant virus endogenization events correlate with presence of 5mC in eukaryotes. Cladogram representing the lineages where large Giant Virus Endogenization Events (GEVEs) have been described in the literature, 5mC and Gene body Methylation pattern (GbM) presence. Dark dots indicate presence, white dots absence and brown dots indicate lack of data/studies. Schematic of the lateral gene transfers (LGT) from giant viruses to eukaryotes, and vice versa, mediated by chromatin regulatory mechanisms. DNA methyltransferases (DNMTs) mediate DNA methyl marks shown as lollipops.

Giant viruses emerged before the origins of modern eukaryotes(16), and chromatin silencing mechanisms such as 5mC or histone modifications were present in the Last Eukaryotic Common Ancestor(1, 9, 45). Thus, these patterns of frequent giant virus endogenization that we observe in modern eukaryotes must have been constant during the whole history of the lineage. Although domestication of giant virus derived genes might be rare, we can see examples of this occurring throughout the tree of life(13, 70, 71). It is worth highlighting that despite giant virus derived genes being widespread, their domestication potential is harder to assess, given the difficulty to test their roles and expression across divergent protist species. Thus, giant viruses, whose genetic material is itself a composite of various origins(72–74), serve as a source of genetic novelty via lateral gene transfer across eukaryotes (**Fig. 5**). Unlike plasmids or other forms of bacterial lateral gene transfer mechanisms, giant viruses are a dangerous vessel for genetic interchange, thus chromatin silencing mechanisms are probably required for a stepwise acquisition of foreign DNA. In turn, the host chromatin-protection is likely counteracted by giant viruses, as exemplified by the histone demethylases present in *Amoebidium* pandoraviruses, or other examples of chromatin modifiers reported in giant virus genomes(45). Similarly, presence of DNMTs in pandoraviruses, and the capacity of giant viruses to modify their own DNA(44), could be a protective response against eukaryotic chromatin, avoiding viral DNA to be recognised as a threat. Chromatin hijacking by giant viruses is a process reminiscent of cases in which TEs have co-opted host chromatin regulators(42, 55, 75, 76), highlighting the age-long conflict between eukaryotic chromatin and parasitic DNA. In summary, *Amoebidium* exemplifies the intricate network-like origins of eukaryotic DNA, challenging traditional notions of strict vertical inheritance within the clade.

## Materials and Methods

### Cell culture, treatment and nucleic acid extraction

*Amoebidium* isolates were grown on Brain Heart Infusion (10% BHI, ThermoFisher CM1135) liquid media at 25°C in 25 mL culture flasks. *Sphaeroforma arctica, Creolimax fragrantissima*, and *Chromosphaera perkinsii* were grown in liquid Marine Broth (Difco™ Marine Broth 2216) at 17°C. Six *Amoebidium* alternative isolates were obtained from the ARS Collection of Entomopathogenic Fungal Cultures.

DNA methylation drugs 5-Azacytidine (ab142744), decitabine (ab120842), and zebularine (ab141264) were dissolved in DMSO. *A. appalachense* was grown with 0M, 0.1µM, 1µM, 10µM, 100µM, and 1mM final concentration of each drug in 2ml 10% BHI 10% DMSO media in a 12 well plate, and effects were tracked daily for 5 days. Only 100µM and 1mM 5-Azacytidine showed a growth phenotype. *A. appalachense* DNA and RNA were extracted from cultures grown for 3 days in 10ml 10% BHI with 1% DMSO, and 1% DMSO with 100µM of their respective drug, in triplicate. 12nmol of 5-Azacytidine in 120ul DMSO was spread over 12ml agar plates of BHI (*A. appalachensis*) and Marine broth (*S. arctica*, *C. fragrantissima*, *C. perkinsii*), and dilution assays for growth were done for all 4 ichthyosporean species using 1x, 10x, 100x, 1,000x and 10,000x serial dilutions of saturated culture.

The developmental cell cycle of *A. appalachensis* was determined using a combination of live and fixed-cell microscopy using a fully motorized Nikon Ti2-E epifluorescence inverted microscope equipped with a hardware autofocus PFS4 system, a Lumencor SOLA SMII illumination system and a Hamamatsu ORCA-spark Digital CMOS camera. A CFI Plan Fluor 20X, 0.50 NA., a CFI Plan Fluor 40X Air and a CFI Plan Fluor 60X Oil, 0.5-1.25 NA. objectives were used for imaging. For live-cell microscopy, a 25 day old culture was diluted 1/250 and imaged with BF every 15 minutes for 72 h at a controlled temperature of 23°C in 600 µl wells using a cooling/heating P Lab-Tek S1 insert (Pecon GmbH) with Lauda Loop 100 circulating water bath. We examined 120 videos counting events of spontaneous cell death, cellularisation, and cell release, and the number of released spores per colony (total 703 cells tracked, Supplementary Table S1). For fluorescent microscopy, samples were fixed in 4% formaldehyde, washed with PBS, and stained with Phalloidin and Hoechst to visualise and count actin and nuclei, respectively every 4-5 hours over 72 hours. RNA and DNA were obtained for representative stages of the life cycle: 5 hours post inoculation (unicell - uninucleated), 14 h (coenocyte), 20h (cellularisation), 33h (cell release).

DNA for *A. appalachense* genome sequencing was extracted using liquid nitrogen grinding and Qiagen MagAttract HMW DNA Kit & QIAGEN Genomic-tip 20/G (10223), and for *A. appalachense*, *S. arctica*, *C. perkinsii*, and *C. fragrantissima* Enzymatic Methyl-seq samples we used NEB Monarch Genomic DNA Purification Kit. DNA for *Amoebidium* isolates 9181 and 9257 was extracted with phenol chloroform extraction and further purification with NEB Monarch Genomic DNA Purification Kit. RNA for all samples was extracted using nitrogen grinding and Monarch Total RNA Miniprep Kit.

### Micro-C library preparation

A. *appalachense* cells grown for 7 days were crosslinked for 10 min with 1% formaldehyde under vacuum conditions in a desiccator. The reaction was quenched with 128 mM glycine for 5 min under vacuum, followed by an additional incubation on ice for 15 min. Crosslinked cells were washed twice and subsequently resuspended in a 1/10th PBS solution. Coenocytic cell walls were disrupted by glass bead beating for 5 min followed by a second crosslinking step with 3 mM DSG for 40 min at room temperature.

Micro-C libraries were prepared as described(77) with the following modifications. In-nuclei chromatin digestion to achieve 80 % monomer / 20 % oligomers nucleosome ratio was performed with MNase (Takara Bio, 2910a) 100 U per 4M nuclei for 10 min. The digested chromatin ends were repaired and labelled with biotinylated nucleotides. Prior to proximity ligation, the digested chromatin was released from nuclei and permeabilized coenocytes by glass bead beating for 10 min. Next, proximal nucleosomes were ligated together, and unligated ends were treated with Exonuclease III (NEB, M0206) to remove biotin-dNTPs. The chromatin was then decrosslinked and deproteinazed, and ligated DNA fragments were captured with Dynabeads MyOne Streptavidin (Life technologies, 65602). Libraries were barcoded using the NEBNext End repair/dA-tailing mix (NEB, E7546) and NEBNext Ultra II Ligation Module (NEB, E7595S). The final amplified libraries, comprising three biological replicates, were sequenced with NextSeq500 in paired-end format with a read length 42 bases per mate, obtaining a total of 131,547,803 sequenced reads.

### Genome sequencing and assembly

High molecular weight genomic DNA from *A. appalachense* was ligated with the Nanopore SQK-LSK110 ligation kit and sequenced in Promethion R9 flowcells. Since pore clogging occurred quickly, we performed short sequencing runs followed by flowcell cleanup steps, and re-loading of fresh library in intervals, requiring 3 flowcells. In parallel, a library of paired-end short-reads generated with the TruSeq kit and sequenced with an Illumina HiSeq2500. Nanopore reads were basecalled using the “sup” model with Guppy (v6.2.1), and assembled with Flye (v2.9-b1768) with the “--nanopore_hq” parameter and two rounds of polishing(78). The resulting genome was further polished with the short-reads with Pilon(79) for two rounds, using BUSCO score (-m genome, v5)(80) to validate improvements, obtaining a contig level N50 of 1.8 Mb. Micro-C data were mapped on the genome using Juicer (v1.6)(81) with the -p assembly option. The 3D-DNA pipeline(82), utilising the proximity ligation data, was used to scaffold the genome with -r3 -editor-repeat-coverage 10. Final manual curation in the Juicebox Assembly Tool(83) resulted in 18 chromosomes. The genome was then polished using Medaka with the original nanopore reads.

For the Isolates 9181 and 9257, we ligated the DNA using Nanopore SQK-LSK114 ligation kit and sequenced following the same strategy but using PromethIon R10 flowcells (Supplementary Table S2). Contig-level assembly was obtained using Flye with Guppy “sup” base called reads as above. Medaka polishing was discarded as it decreased BUSCO score. Then D-GENIES was used to visualise the synteny with the reference genome(84). RagTag scaffolding using the reference genome was performed for both Isolate contigs(85), yet only 9181 was kept as 98% of the sequence were placed into chromosomes, whereas 9257 only got 54%, rendering the scaffolding unreliable. Extra scaffolding using P_RNA_scaffolder(86) was performed for 9257 using its transcriptomic data to further increase contiguity, and validated through BUSCO improvement criteria.

### Genome annotation

We generated a *de novo* RepeatModeler2(87) annotation with the LTR module to characterise *A. appalachense* repeat landscape. This was then mapped to the genome using RepeatMasker. In parallel, publicly available deep coverage RNA-seq from *A. appalachense* (SRR545192) was mapped to the genome using HISAT2 with the –dta parameter, and Stringtie for reference based transcriptome assembly(88). The resulting bam was processed with Portcullis to generate a list of high-quality intron junctions(89). In parallel, *de novo* Trinity assembly of the SRR545192 reads was mapped using gmap to the genome(90). The combination of introns, Stringtie and Trinity mappings was fed to Mikado to choose the best collection of transcripts based on the Uniprot Sprot database. The best transcripts were used to train Augustus model for *Amoebidium*(91). To inform Augustus annotation, we mapped protein alignments against the genome using MetaEuk(92), using closely related high-quality ichthyosporean genomes as query, obtaining coding sequence hints. Portcullis introns and Mikado exons were also introduced as hints for Augustus genome annotation. The resulting Augustus annotation was then updated using PASA with the Mikado transcripts, fixing broken gene models and adding UTRs. Annotation was visually inspected in the IGV genome browser. Functional annotation was obtained using hmmscan with Pfam-A database(93) and the eggNOG-mapper server(94).

To annotate the alternative Isolate genomes, the same process was followed, using the reference annotation for the MetaEuk CDS hints, and the pre-trained *Amoebidium* Augustus model. All annotations were evaluated using BUSCO v5 with eukaryota_odb10 database.

### Transcriptome sequencing, assembly and analysis

We used 50 to 1000 ng of RNA from treated samples, developmental time points and Isolates to build mRNA-seq libraries (see Supplementary Table S3 for details), first enriching for poly-A transcripts with the NEB Magnetic mRNA Isolation Kit S1550S, and then building the libraries with the NEBNext® Ultra™ II Directional RNA Library Prep Kit for Illumina® (E7760L) according to manufacturer’s instructions. Short read Illumina reads were obtained with a NovaSeq6000. De novo transcriptome assemblies were obtained with Trinity (strand-specific) for the Isolates. The Trinity assemblies were searched for 18S and 16S sequences using BLASTn with NCBI query sequences.

Drug-treatment and developmental samples were mapped against the annotation using Kallisto to obtain TPMs(95). To perform differential expression analysis of TEs and protein-coding genes, we used HISAT2 with the TElocal pipeline(96), obtaining gene counts that were then analysed in DEseq2(97). Only intergenic TEs above 500 bp were kept for the analysis.

*Sphaeroforma* treatment samples were done in the same way and mapped to the latest version of the genome(98).

### Methylome sequencing and analysis

We sonicated genomic DNA from *A. appalachense* (control, developmental timepoints, DMSO / 5-Azacytidine treated), *S. arctica, C. perkinsii* and *C. fragrantissima*, spiked with phage lambda DNA and methylated pUC19 controls, to obtain 300 bp fragments with a Covaris M220. Then, we used the NEB Enzymatic Methyl-Kit to convert all the unmethylated Cs into Ts as described in the manufacturers instructions(99). These libraries were then sequenced in a Illumina NovaSeq6000 to various coverages (Supplementary Table S4). The reads were then mapped with fastp and mapped to the reference genomes (27, 98, 100) using BS-Seeker2 backed with bowtie2(101). Sambamba was used to remove PCR duplicates and CgmapTools was used to obtain the methyl calls(102). These files were processed in R using the bsseq package, and bigwig tracks using the BedGraphToBigWig UCSC utility.

In parallel, Nanopore reads were basecalled and mapped for base modifications using the Guppy dna_r9.4.1_450bps_modbases_5mc_cg_sup_prom.cfg and dna_r10.4.1_e8.2_400bps_modbases_5mc_cg_sup_prom.cfg models. The resulting read alignments were processed with modbam2bed the --cpg -e -m 5mC parameters. These bed files were also processed in R using the bsseq package.

### Giant virus identification and phylogenetic analysis

Visual inspection of hypermethylated blocks revealed core giant virus genes in unusual gene architecture patterns. To validate these potential claims, we used ViralRecall(103) that flagged just a few of these sequences as potential giant virus endogenization events. However, we observed that many events were not captured by that software, so we manually inspected the genome to obtain the longest potential inserts, filtering out TEs inserted within the viral region. We searched those consensus sequences against the genome using BLASTn, to obtain all potential regions of homology to giant viruses. Another round of manual inspection of all chromosomes using non-CGC/GCG methylation blocks as boundary demarcation was used to delimit integration sites. The same process was used for Adintoviruses and Plavaka giant repeats.

Hmmsearch was used to identify core viral genes, DNMTs (PF00145) and JmjC (PF02373) containing proteins. The obtained genes were included to reference databases (ref giant virus DB, adintovirus, Eukaryotic chromatin NEE, DNMTs) and aligned using MAFFT in lins-i mode(104). Alignments were trimmed using TrimAL with the -gappyout mode(105). The resulting alignments were fed into IQ-TREE 2 with automatic model testing to build maximum likelihood phylogenetic trees using altr and uboot as nodal support measures(106). Adintovirus minor and major capsid proteins were annotated with HHpred against PDB_mmCIF70 database.

Comparative genomics among giant virus insertions (used as independent taxa) or across the Isolate genomes was performed using OrthoFinder with DIAMOND as a search engine(107).

## Acknowledgements

We thank Chema Martín-Durán, Sam Buckberry and Ferdinand Marletaz for reading and giving feedback on this manuscript. We thank Meritxell Antó and Iñaki Ruiz-Trillo for sharing the culture and DNA samples, and Atsushi Toyoda at National Institute of Genetics in Japan for technical support for NGS sequencing. We thank Billie E. Davies for help with Nanopore library construction, and the technical staff at the Department of Biology at Queen Mary University of London for their support, specially William Tyne and Giulia Mastroianni. This work used computing resources from Queen Mary University of London’s Apocrita HPC facilities. This work was funded by the Horizon 2020 Framework Programme to AdM (European Research Council Starting Grant action number 950230), LS was funded by a QMUL PhD fellowship, IVK was supported by Juan de la Cierva postdoctoral fellowship (FJC2020-043131-I) from Spanish Ministry of Science and Innovation, HS was supported by JSPS KAKENHI Grant Number JP16H06279 (PAGS) and JP26891021, OD and MO were funded by an Ambizione fellowship from the Swiss National Science Foundation (PZ00P3_185859), research in ASP group was supported by the European Research Council (ERC-StG 851647) and the Spanish Ministry of Science and Innovation grant (PID2021-124757NB-I00) as well the Spanish Ministry’s support to the EMBL partnership, the Centro de Excelencia Severo Ochoa and the CERCA Programme (Generalitat de Catalunya).

## Contributions

AdM conceived and designed the study. LS performed all experiments including nanopore sequencing, EM-seq, RNA-seq, and drug treatments. LS, AdM and VO performed bioinformatic analysis. IK and ASP performed Micro-C and genome scaffolding. HS provided Illumina sequencing data. MO, OD and LS performed growth, imaging and drug experiments. AdM and LS drafted the manuscript, and all authors critically read and commented on the manuscript.

