## SupplementaryFigures for "DNA methylation enables recurrent endogenization of giant viruses in an animal relative"

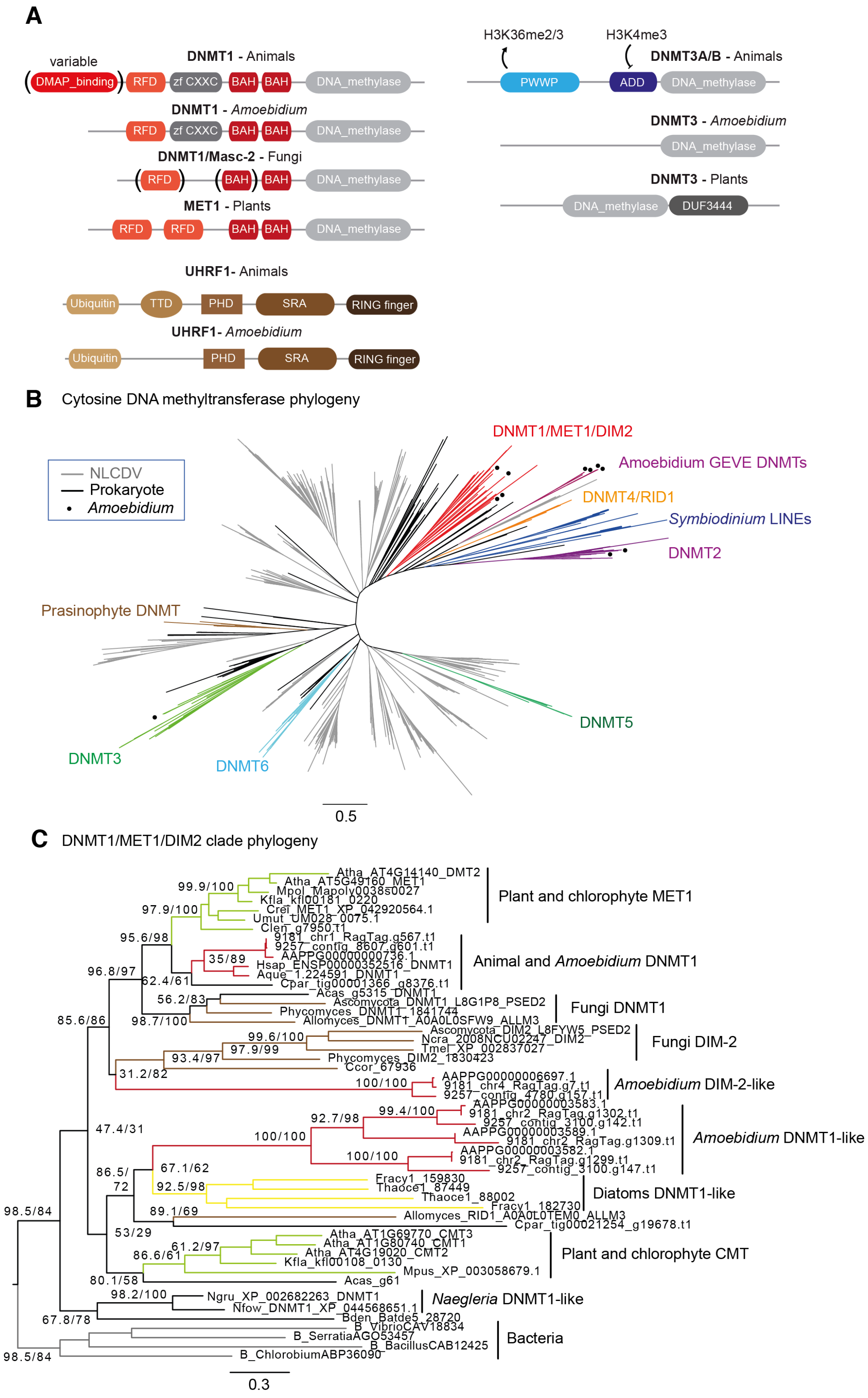
A
variable
DNMT1 - Animals
DMAP_binding
RFD
zf CXXC
BAH
BAH
DNA_methylase
DNMT1 - Amoebidium
DNMT1/Masc-2 - Fungi
MET1 - Plants
UHRF1- Animals
Ubiquitin
TTD
PHD
SRA
RING finger
UHRF1- Amoebidium
H3K36me2/3
H3K4me3
DNMT3A/B - Animals
PWWP
ADD
DNMT3 - Amoebidium
DNMT3 - Plants
DUF3444
B
Cytosine DNA methyltransferase phylogeny
NLCDV
Prokaryote
Amoebidium
DNMT1/MET1/DIM2
Amoebidium GEVE DNMTs
DNMT4/RID1
Symbiodinium LINEs
DNMT2
Prasinophyte DNMT
DNMT5
DNMT3
DNMT6
0.5
C
DNMT1/MET1/DIM2 clade phylogeny
Atha_AT4G14140_DMT2
Atha_AT5G49160_MET1
Mpol_Mapoly0038s0027
Kfla_kfl00181_0220
Crei_MET1_XP_042920564.1
Umut_UM028_0075.1
Clen_g7950.t1
9181_chr1_RagTag.g567.t1
9257_contig_8607.g601.t1
AAPPG00000000736.1
Hsap_ENSP00000352516_DNMT1
Aque_1.224591_DNMT1
Cpar_tig00001366_g8376.t1
Acas_g5315_DNMT1
Ascomycota_DNMT1_L8G1P8_PSED2
Phycomyces_DNMT1_1841744
Allomyces_DNMT1_A0A0L0SFW9_ALLM3
Ascomycota_DIM2_L8FYW5_PSED2
Ncra_2008NCU02247_DIM2
Tmel_XP_002837027
Phycomyces_DIM2_1830423
Ccor_67936
AAPPG00000006697.1
9181_chr4_RagTag.g7.t1
9257_contig_4780.g157.t1
AAPPG00000003583.1
9181_chr2_RagTag.g1302.t1
9257_contig_3100.g142.t1
AAPPG00000003589.1
9181_chr2_RagTag.g1309.t1
AAPPG00000003582.1
9181_chr2_RagTag.g1299.t1
9257_contig_3100.g147.t1
Fracy1_159830
Thaoce1_87449
Thaoce1_88002
Fracy1_182730
Allomyces_RID1_A0A0L0TEM0_ALLM3
Cpar_tig00021254_g19678.t1
Atha_AT1G69770_CMT3
Atha_AT1G80740_CMT1
Atha_AT4G19020_CMT2
Kfla_kfl00108_0130
Mpus_XP_003058679.1
Acas_g61
Ngru_XP_002682263_DNMT1
Nfow_DNMT1_XP_044568651.1
Bden_Batde5_28720
B_VibrioCAV18834
B_SerratiaAGO53457
B_BacillusCAB12425
B_ChlorobiumABP36090
99.9/100
97.9/100
95.6/98
35/89
96.8/97
62.4/61
56.2/83
85.6/86
98.7/100
99.6/100
97.9/99
93.4/97
31.2/82
100/100
99.4/100
92.7/98
47.4/31
100/100
100/100
86.5/
72
67.1/62
92.5/98
89.1/69
98.5/84
53/29
61.2/97
86.6/61
80.1/58
98.2/100
67.8/78
98.5/84
Plant and chlorophyte MET1
Animal and Amoebidium DNMT1
Fungi DNMT1
Fungi DIM-2
Amoebidium DIM-2-like
Amoebidium DNMT1-like
Diatoms DNMT1-like
Plant and chlorophyte CMT
Naegleria DNMT1-like
Bacteria
0.3

**Fig. S1.**

**Distribution of DNMTs in *Amoebidium*.** (**A**) Protein domain architectures of DNMT members of DNMT1 and DNMT3 families as well as the UHRF1, the heterodimer of DNMT1, in various eukaryotic groups. Domains are defined by PFAM: DMAP_binding (PF06464), RFD (PF12047), zf-CXXC (PF02008), BAH (PF01426), DNA_methylase (PF00145), Ubiquitin (PF00240), Tandem Tudor Domain (TDD, PF12148), PHD (PF00628), SRA (PF02182), PWWP (PF00855), ADD (PF17980). Parenthesis indicates variability in the presence of the domain within the group. Arrows indicate that the PWWP domain of DNMT3A/B orthologues in mammals binds to H3K36me2/3, whereas the ADD domain prevents binding to H3K4me3 marked regions. (**B**) Maximum-likelihood phylogenetic tree of DNMTs including prokaryotic, eukaryotic and giant virus sequences. Known eukaryotic groups are depicted with colours, and presence of *Amoebidium* sequences are highlighted with a black dot. (**C**) Maximum-likelihood phylogeny of the clade of DNMT1 and DNMT1-like sequences in eukaryotes, rooted with the closest bacterial sequences. Nodal supports display ALRT and uboot values calculated by IQTREE. The bottom bar indicates substitutions per site. Species IDs: Atha (*Arabidopsis thaliana*), Mpol (*Marchantia polymorpha*), Kfla (*Klebsormidium nitens*), Crei (*Chlamydomonas reinhardtii*), Umut (*Ulva mutabilis*), Clen (*Caulerpa lentillifera*), Hsap (*Homo sapiens*), Aque (*Amphimedon queenslandica*), Cpar (*Cyanophora paradoxa*), Acas (*Acanthamoeba castellannii*), Ncra (*Neurospora crassa*),Tmel (*Tuber melanosporum*), Ccor (*Conidiobolus coronatus)*, Fracy (*Fragilariopsis cylindrus*), Thaoce (*Thalassiosira oceanica*), Mpus (*Micromonas pusilla*), Ngru (*Naegleria gruberi*), Nfow (*Naegleria fowleri*), APARG/9181/9257 (*Amoeibidium* and isolates).

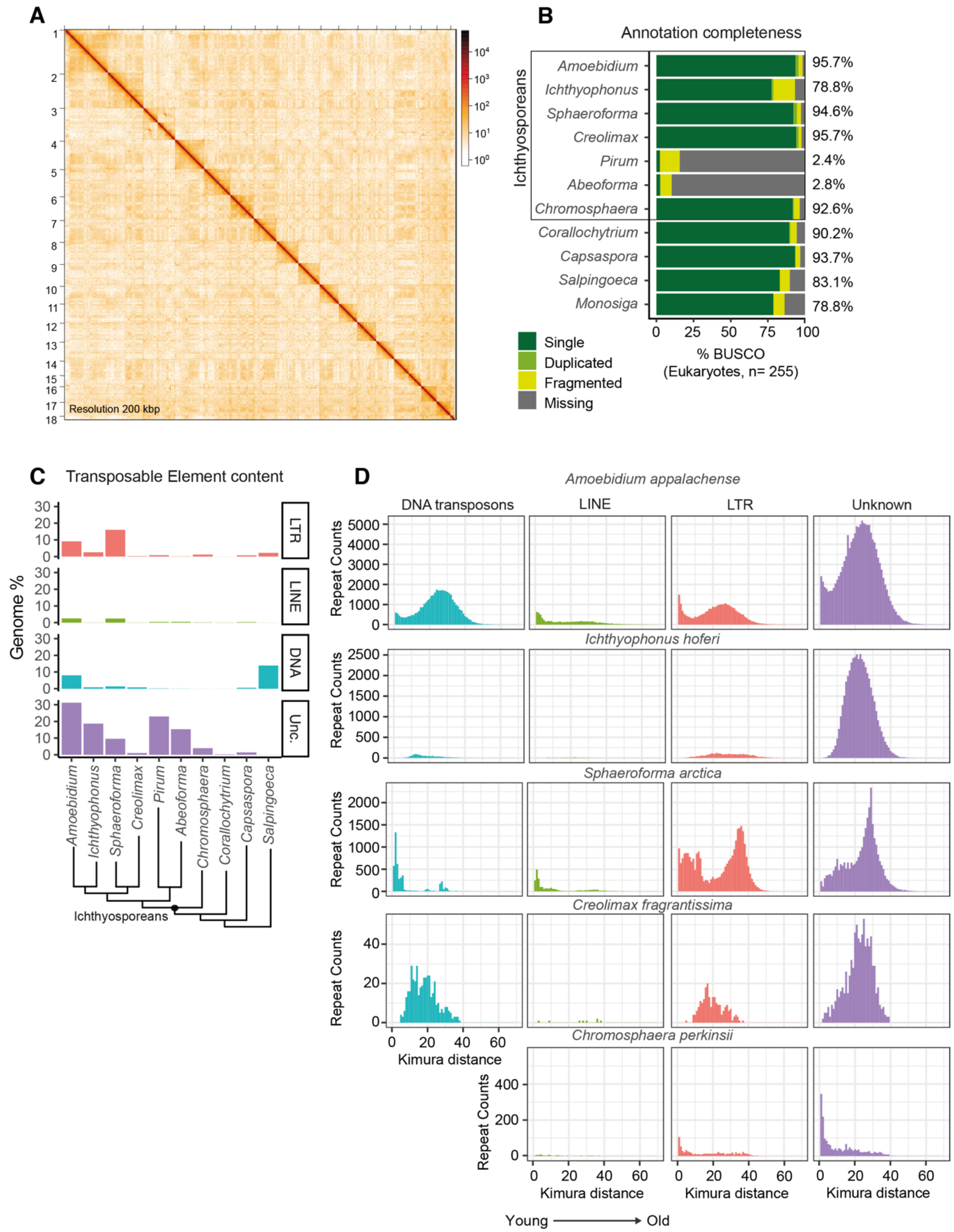


**Fig. S2.**

**Genome assembly, completeness and repeat landscape in *Amoebidium*.** (**A**) Heatmap representing the micro-C contact map across and within *Amoebidium* chromosomes, with a window resolution size of 200 kb. (**B**) BUSCO comparison across published unicellular holozoan genomes, with the total BUSCO % (Single + Duplicated %) displayed on the right hand side. (**C**) Major TE type contributions to the genomes of selected unicellular holozoans. TEs annotations have been obtained with RepeatModeler2 and RepeatMasker. LTR stands for long terminal repeat retrotransposons, LINE for non-LTR retrotransposons, DNA for DNA transposons and Unc for Unknown / unclassified repeats. (**D**) Age profile of repeats across high quality ichthyosporean genomes obtained as Kimura distances (not corrected for CpGs).

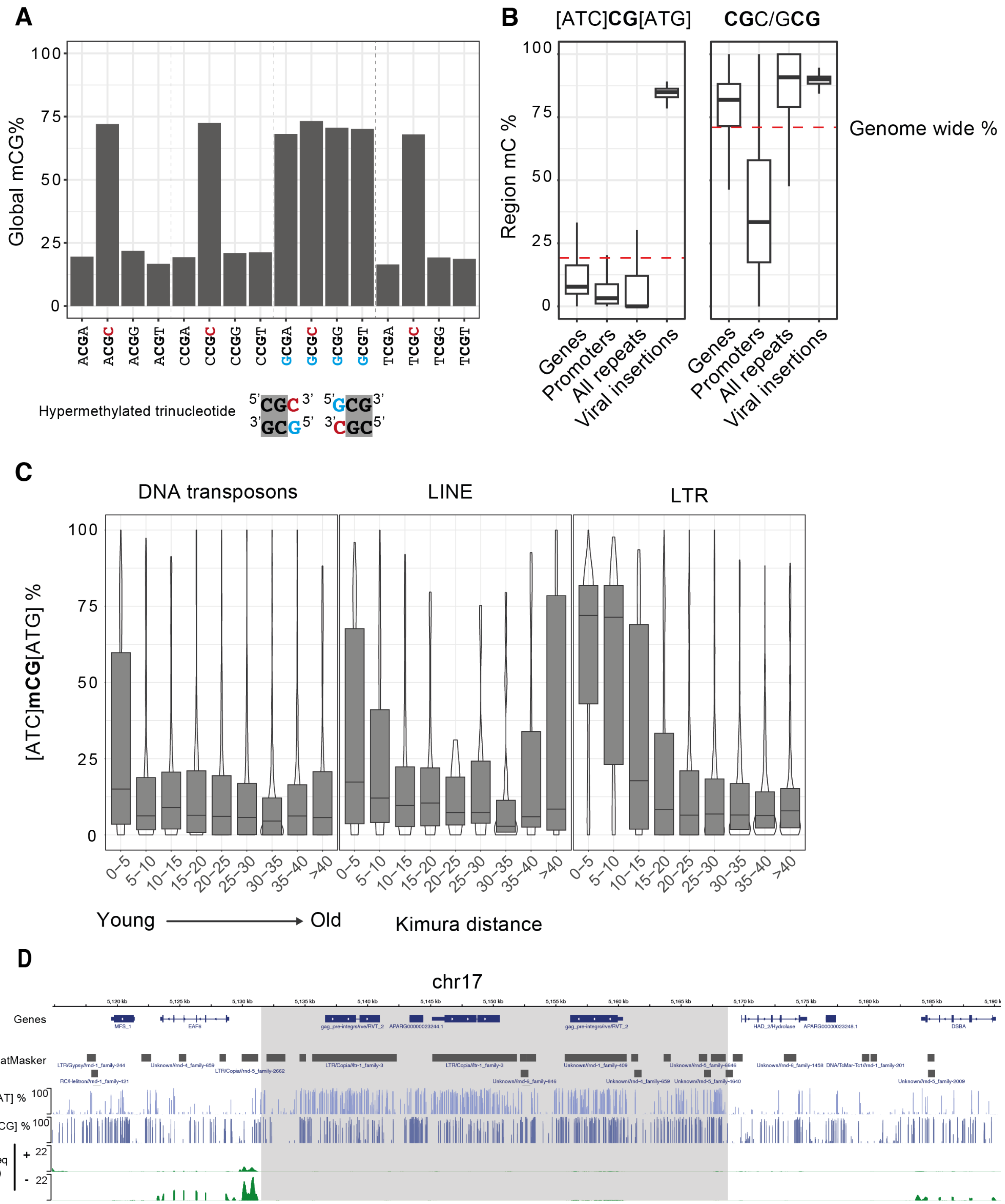


**Fig. S3.**

**Trinucleotide context influences CG methylation distribution in *Amoebidium*.** (**A**) Genome wide methylation levels on CG dinucleotides surrounded by all possible base compositions measured by Enzymatic Methyl-seq. (**B**) Regional methylation across genes, promoters, repeats and viral insertions divided by trinucleotide context. (**C**) non-CGC/GCG methylation distribution on TEs classified by age. TE insertions with Kimura distances spanning 5% differences are shown from left to right. Centre lines in boxplots are the median, box is the interquartile range

(IQR), and whiskers are the first or third quartile ± 1.5× IQR, outliers not shown. (**D**) Genome browser snapshot of retrotransposons methylated in all CG contexts, and neighbouring genes predominantly displaying CGC/GCG methylation.

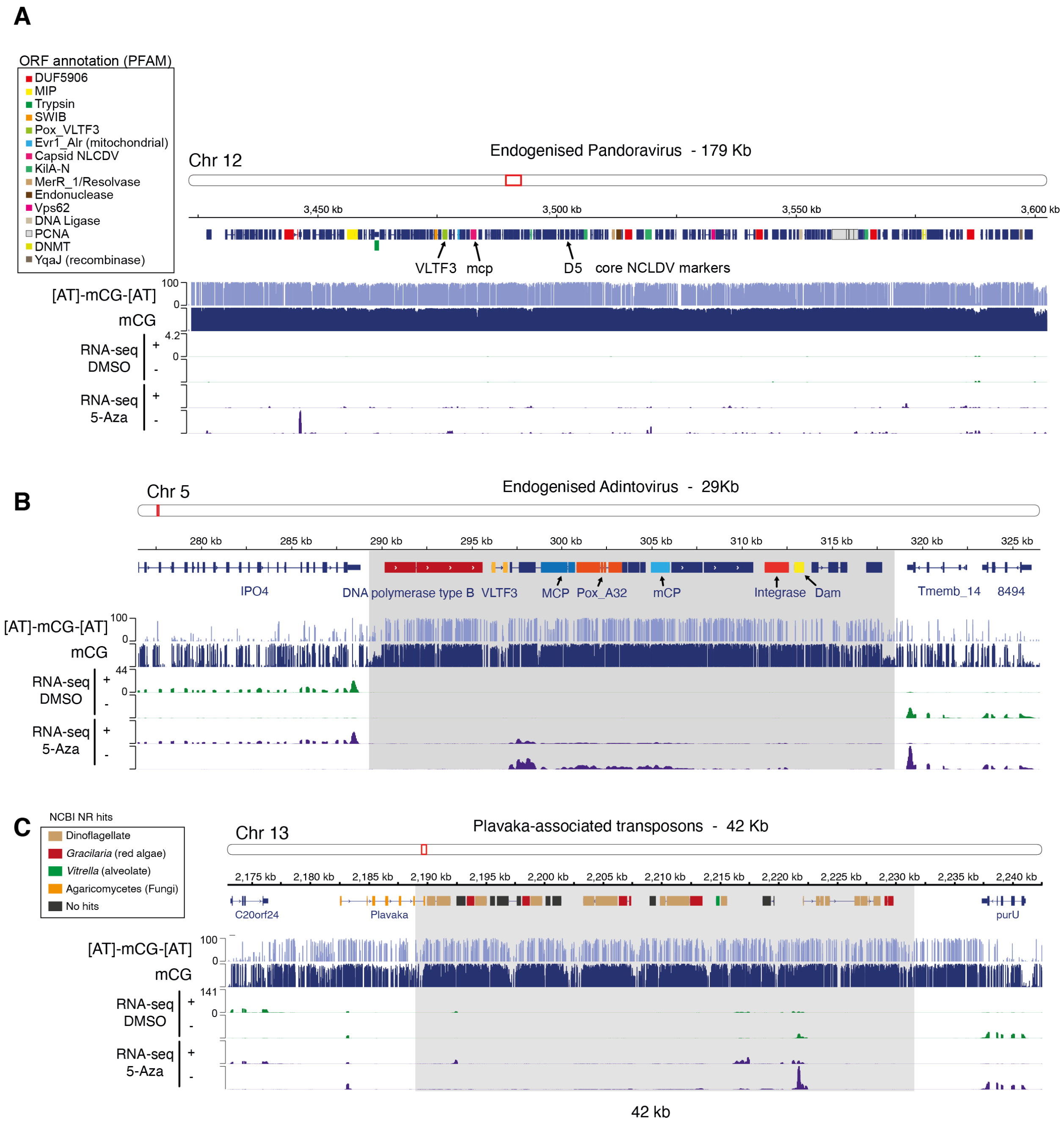


**Fig. S4.**

**The three major giant repeats in the *Amoebidium* genome.** (**A**) Genome browser snapshot of a large pandoravirus insertion in chromosome 12. Coloured genes highlight various PFAM containing Open Reading Frames. (**B**) Genome snapshot of an Adintovirus insertion spanning 29 kb, flanked by host protein coding genes. mCP: minor capsid protein, MCP: major capsid protein. (**C**) Plavaka-associated giant repeat spanning 42 kb. Genes are colour coded according to the taxonomy of the first hits against the NCBI non-redundant protein database.

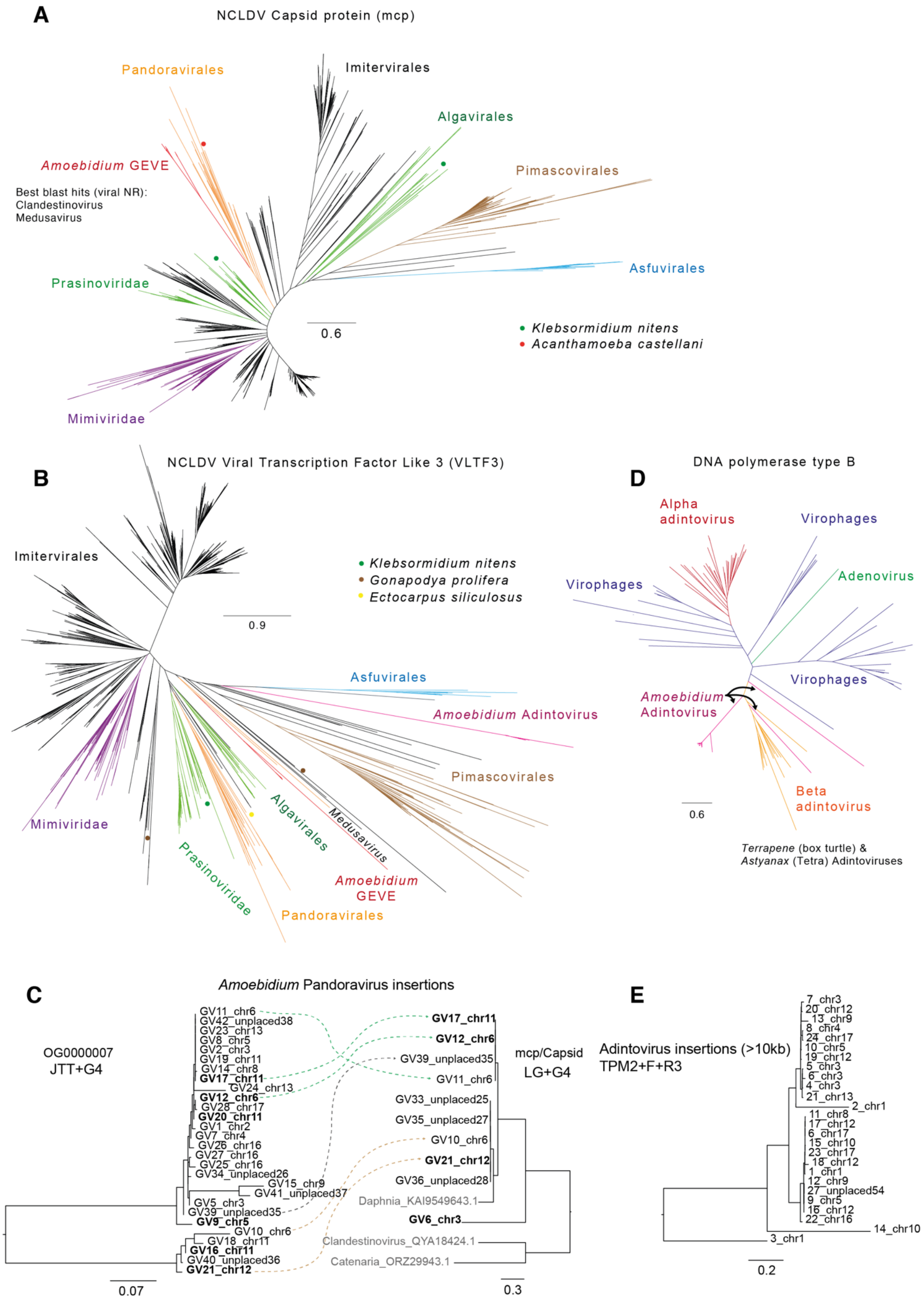
A
NCLDV Capsid protein (mcp)
Pandoravirales
Imitervirales
Algavirales
Amoebidium GEVE
Pimascovirales
Best blast hits (viral NR):
Clandestinovirus
Medusavirus
Asfuvirales
Prasinoviridae
0.6
Klebsormidium nitens
Acanthamoeba castellani
Mimiviridae
B
NCLDV Viral Transcription Factor Like 3 (VLTF3)
Imitervirales
Klebsormidium nitens
Gonapodya prolifera
Ectocarpus siliculosus
0.9
Asfuvirales
Amoebidium Adintovirus
Pimascovirales
Mimiviridae
Algavirales
Medusavirus
Prasinoviridae
Amoebidium
GEVE
Pandoravirales
D
DNA polymerase type B
Alpha
adintovirus
Virophages
Virophages
Adenovirus
Virophages
Amoebidium
Adintovirus
Beta
adintovirus
0.6
Terrapene (box turtle) &
Astyanax (Tetra) Adintoviruses
C
Amoebidium Pandoravirus insertions
OG0000007
JTT+G4
mcp/Capsid
LG+G4
GV11_chr6
GV42_unplaced38
GV23_chr13
GV8_chr5
GV2_chr3
GV19_chr11
GV14_chr8
GV17_chr11
GV24_chr13
GV12_chr6
GV28_chr17
GV20_chr11
GV1_chr2
GV7_chr4
GV26_chr16
GV27_chr16
GV25_chr16
GV34_unplaced26
GV15_chr9
GV41_unplaced37
GV5_chr3
GV39_unplaced35
GV9_chr5
GV10_chr6
GV18_chr11
GV16_chr11
GV40_unplaced36
GV21_chr12
GV17_chr11
GV12_chr6
GV39_unplaced35
GV11_chr6
GV33_unplaced25
GV35_unplaced27
GV10_chr6
GV21_chr12
GV36_unplaced28
Daphnia_KAI9549643.1
GV6_chr3
Clandestinovirus_QYA18424.1
Catenaria_ORZ29943.1
0.07
0.3
E
Adintovirus insertions (>10kb)
TPM2+F+R3
7_chr3
20_chr12
13_chr9
8_chr4
24_chr17
10_chr5
19_chr12
5_chr3
6_chr3
4_chr3
21_chr13
2_chr1
11_chr8
17_chr12
6_chr17
15_chr10
23_chr17
18_chr12
1_chr1
12_chr9
27_unplaced54
9_chr5
16_chr12
22_chr16
14_chr10
3_chr1
0.2

**Fig. S5.**

**Phylogenetic alliances of *Amoebidium* endogenized viruses.** (**A**) Maximum-likelihood phylogeny of the NCLDV major capsid protein (PF04451) found in the Giant Virus Database, including select GEVEs. *Amoebidium* sequences are found in the red clade embedded in the pandoravirales clade. Potentially endogenized capsid proteins in the algae *Klebsormidium nitens* and the amoebozoan *Acanthamoeba castellanni* displayed with coloured dots. (**B**) Maximum-likelihood phylogeny of the VLTF3 (Pox_VLTF3, PF04947) genes found in the Giant Virus database and select eukaryotes. *Amoebidium* pandoravirus GEVE are shown in a red branch sister group to pandoravirales, including Medusavirus. *Amoebidium* adintoviruses are shown in purple, no other polinton-like viruses are shown as these do not normally encode VLTF3, which was probably acquired from a giant virus. (**C**) Amino acid maximum-likelihood phylogeny of a widely distributed Orthogroup (OG000007) across Pandoravirus insertions, and major capsid proteins with outgroups from panel A. Each pandoravirus insertion is named as GV[number], and their chromosome of origin. Dotted lines connect insertions that encode both ORFs. (**D**) Maximum-likelihood phylogeny of DNA polymerase type B ORF from adintoviruses, adenoviruses and virophages from (39). (**E**) Nucleotide maximum-likelihood phylogeny of full-length adintovirus insertions. These are labelled with numbers reflecting chromosome of origin. Bars depict the substitutions per site in all phylogenetic trees.

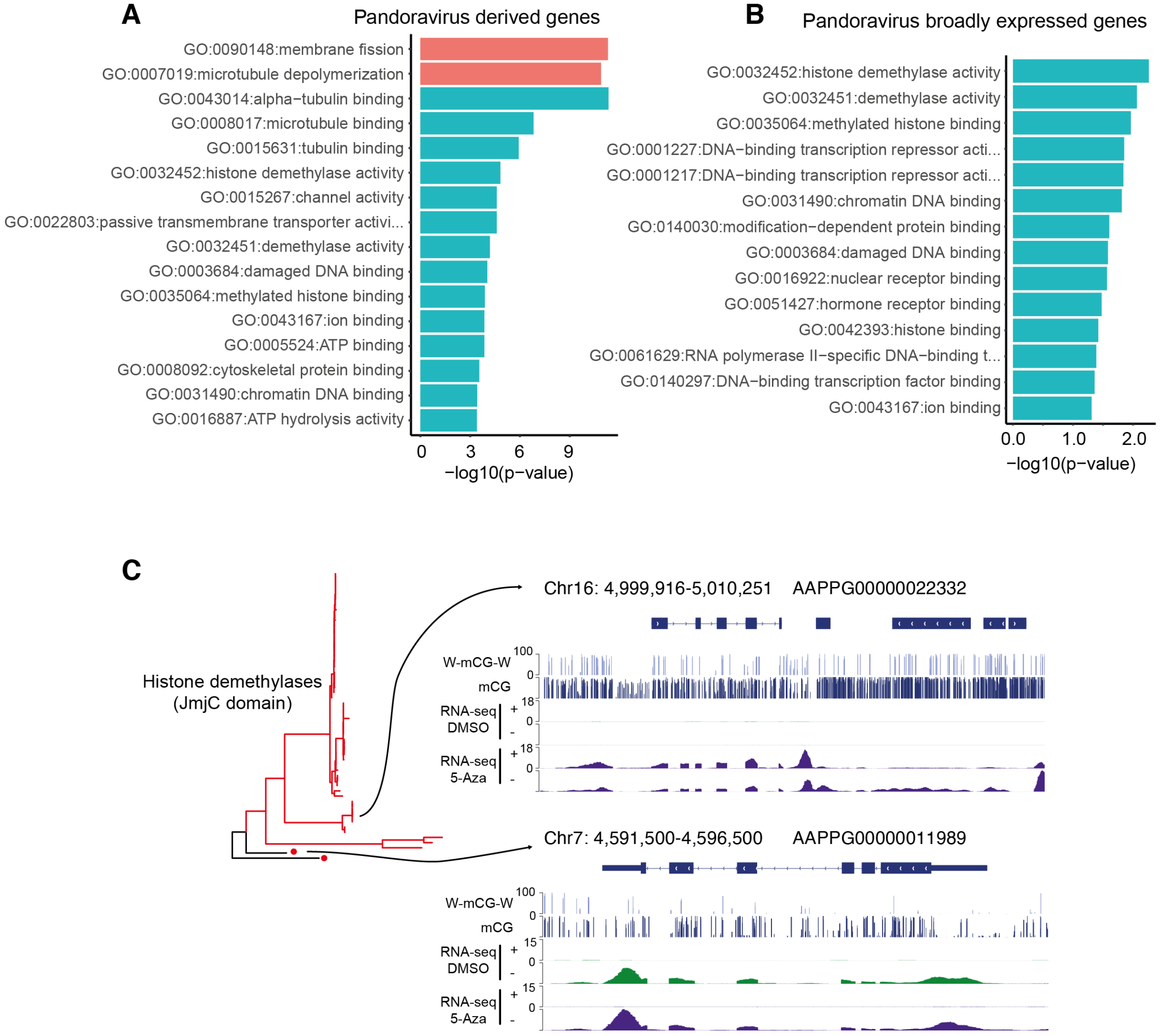


**Fig. S6.**

**Pandoravirus-derived genes are enriched for histone-demethylase activity.** (**A**) Gene Ontology enrichment for all genes encoded in pandoravirus insertions with the background of all *Amoebidium* genes. Red bars indicate “Biological Process” GO categories, while turquoise bars indicate “Molecular Function” GOs. (**B**) Gene ontology enrichment of genes encoded in pandoravirus insertions that are transcribed in culture conditions (TPM >1). (**C**) Genome browser snapshots of KDM4-like genes found in Pandoravirus insertions (chr16, high [ATC]mCG[ATG] methylation) or found as host genes (chr 4, lack of [ATC]mCG[ATG] methylation). RNA-seq tracks are divided per strand (positive or negative) and display control and 5-Azacytidine treatment conditions described in **Fig. 3A**.

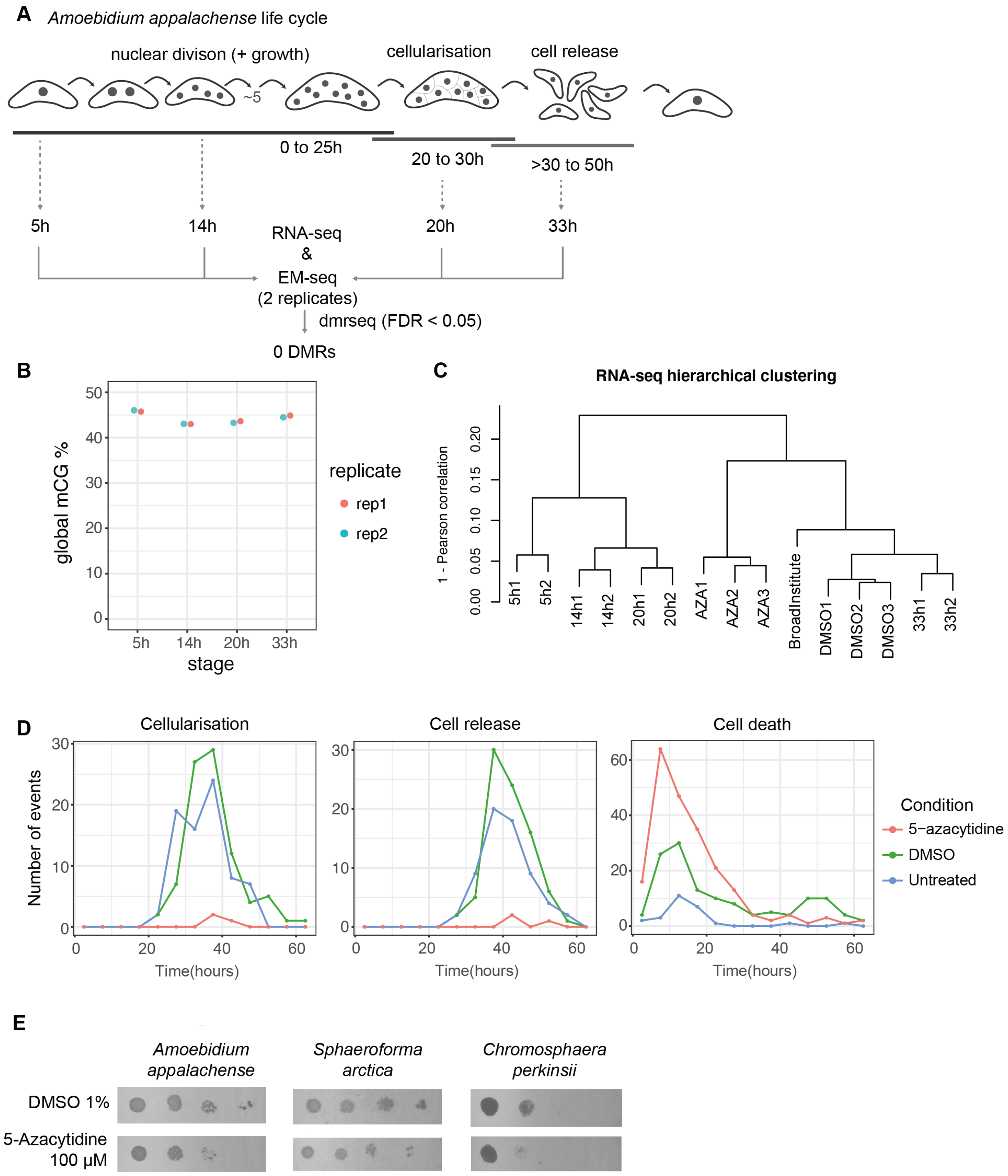

A
Amoebidium appalachense life cycle
nuclear divison (+ growth)
~5
cellularisation
cell release
0 to 25h
20 to 30h
>30 to 50h
5h
14h
20h
33h
RNA-seq
&
EM-seq
(2 replicates)
dmrseq (FDR < 0.05)
0 DMRs
B
global mCG %
stage
replicate
rep1
rep2
C
RNA-seq hierarchical clustering
1 - Pearson correlation
5h1
5h2
14h1
14h2
20h1
20h2
AZA1
AZA2
AZA3
BroadInstitute
DMSO1
DMSO2
DMSO3
33h1
33h2
D
Cellularisation
Cell release
Cell death
Number of events
Time(hours)
Condition
5−azacytidine
DMSO
Untreated
E
Amoebidium appalachense
Sphaeroforma arctica
Chromosphaera perkinsii
DMSO 1%
5-Azacytidine 100 μM


**Fig. S7.**

***Amoebidium* developmental methylation dynamics and response to 5-Azacytidine treatment**. (**A**) Scheme showing the average timings of *Amoebidium appalachense* coenocytic life cycle. Nuclear divisions within a coenocyte last for about 25 hours, reaching 16 to 32 nuclei. Cellularization happens around 20h post culture start, and cellular release past 30h. Sampled time points for RNA-seq and Enzymatic Methyl-seq are shown. The R package dmrseq was used to identify differentially methylated regions (DMRs) across developmental stages, finding 0 DMRs passing the non-conservative FDR threshold of 0.05. (**B**) Global CG methylation levels of developmental Enzymatic Methyl-seq replicates. (**C**) Hierarchical clustering of RNA-seq samples of developmental and control (DMSO)/5-Azacitidine (AZA) treatments clustered by Pearson correlation distance calculated in R. (**D**) Distribution of cellularisation, cellular release and spontaneous death events recorded in live microscopy videos of *Amoebidium appalachense* cultures grown in 10% BHI media, 10% BHI with DMSO at 1% and 10% BHI with 5-Azacytidine at 100 uM. (**E**) Agar plates showing *Amoebidium* colony growth in 10% BHI or *Sphaeroforma* and *Chromosphaera* in Marine Broth media upon DMSO and 5-Azacytidine treatments. Left to right: 5µl 1x, 10x, 100x, 1,000x and 10,000x serial dilutions of saturated culture.

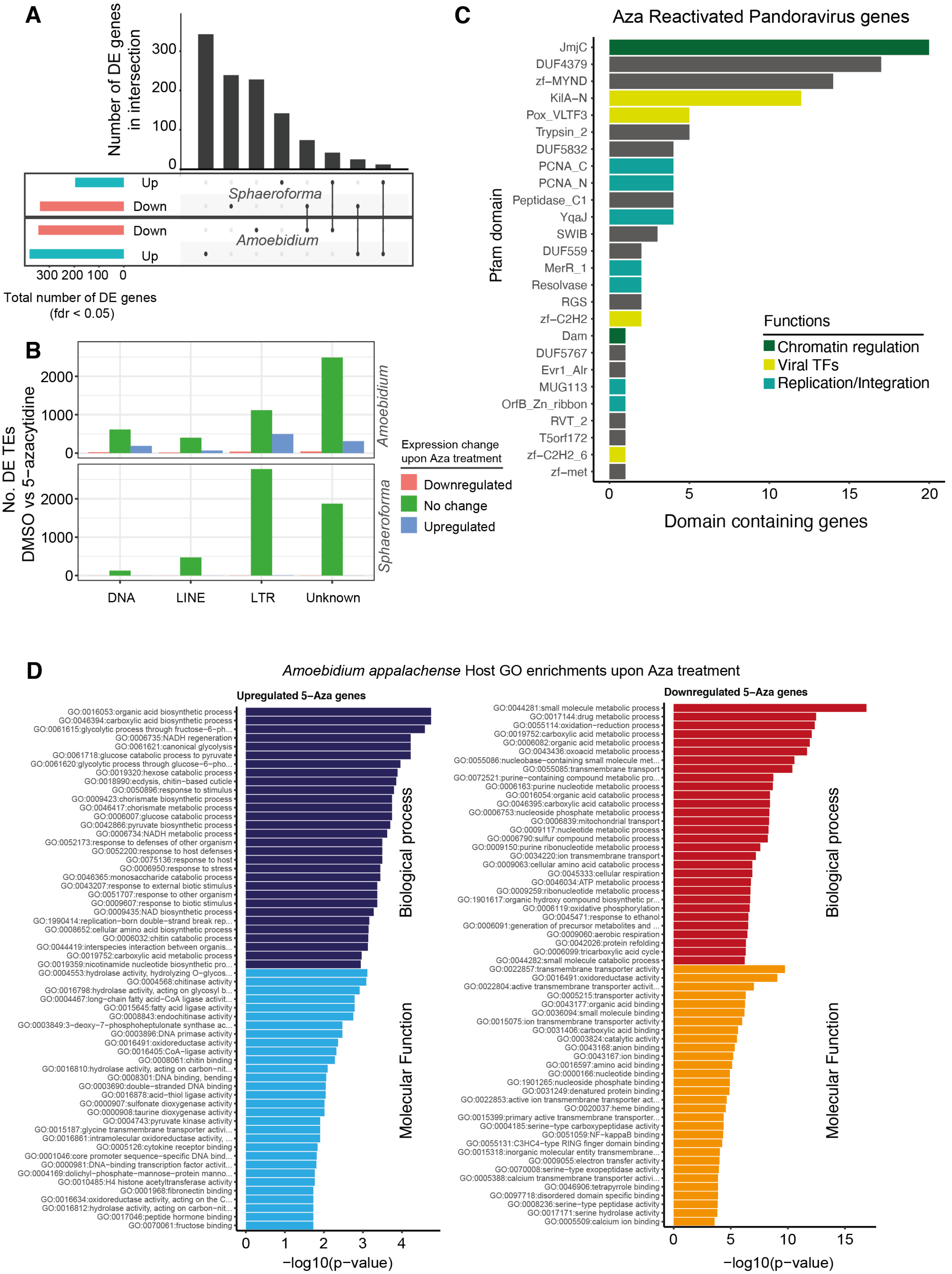

A
Number of DE genes in intersection
300
200
100
0
Up
Down
Sphaeroforma
Down
Amoebidium
Up
300 200 100 0
Total number of DE genes (fdr < 0.05)
B
No. DE TEs DMSO vs 5−azacytidine
2000
1000
0
Amoebidium
Sphaeroforma
DNA
LINE
LTR
Unknown
Expression change upon Aza treatment
Downregulated
No change
Upregulated
C
Aza Reactivated Pandoravirus genes
Pfam domain
JmjC
DUF4379
zf−MYND
KilA−N
Pox_VLTF3
Trypsin_2
DUF5832
PCNA_C
PCNA_N
Peptidase_C1
YqaJ
SWIB
DUF559
MerR_1
Resolvase
RGS
zf−C2H2
Dam
DUF5767
Evr1_Alr
MUG113
OrfB_Zn_ribbon
RVT_2
T5orf172
zf−C2H2_6
zf−met
0
5
10
15
20
Domain containing genes
Functions
Chromatin regulation
Viral TFs
Replication/Integration
D
Amoebidium appalachense Host GO enrichments upon Aza treatment
Upregulated 5−Aza genes
GO:0016053:organic acid biosynthetic process
GO:0046394:carboxylic acid biosynthetic process
GO:0061615:glycolytic process through fructose−6−ph...
GO:0006735:NADH regeneration
GO:0061621:canonical glycolysis
GO:0061718:glucose catabolic process to pyruvate
GO:0061620:glycolytic process through glucose−6−pho...
GO:0019320:hexose catabolic process
GO:0018990:ecdysis, chitin−based cuticle
GO:0050896:response to stimulus
GO:0009423:chorismate biosynthetic process
GO:0046417:chorismate metabolic process
GO:0006007:glucose catabolic process
GO:0042866:pyruvate biosynthetic process
GO:0006734:NADH metabolic process
GO:0052173:response to defenses of other organism
GO:0052200:response to host defenses
GO:0075136:response to host
GO:0006950:response to stress
GO:0046365:monosaccharide catabolic process
GO:0043207:response to external biotic stimulus
GO:0051707:response to other organism
GO:0009607:response to biotic stimulus
GO:0009435:NAD biosynthetic process
GO:1990414:replication−born double−strand break rep...
GO:0008652:cellular amino acid biosynthetic process
GO:0006032:chitin catabolic process
GO:0044419:interspecies interaction between organis...
GO:0019752:carboxylic acid metabolic process
GO:0019359:nicotinamide nucleotide biosynthetic pro...
GO:0004553:hydrolase activity, hydrolyzing O−glycos...
GO:0004568:chitinase activity
GO:0016798:hydrolase activity, acting on glycosyl b...
GO:0004467:long−chain fatty acid−CoA ligase activit...
GO:0015645:fatty acid ligase activity
GO:0008843:endochitinase activity
GO:0003849:3−deoxy−7−phosphoheptulonate synthase ac...
GO:0003896:DNA primase activity
GO:0016491:oxidoreductase activity
GO:0016405:CoA−ligase activity
GO:0008061:chitin binding
GO:0016810:hydrolase activity, acting on carbon−nit...
GO:0008301:DNA binding, bending
GO:0003690:double−stranded DNA binding
GO:0016878:acid−thiol ligase activity
GO:0000907:sulfonate dioxygenase activity
GO:0000908:taurine dioxygenase activity
GO:0004743:pyruvate kinase activity
GO:0015187:glycine transmembrane transporter activi...
GO:0016861:intramolecular oxidoreductase activity, ...
GO:0005126:cytokine receptor binding
GO:0001046:core promoter sequence−specific DNA bind...
GO:0000981:DNA−binding transcription factor activit...
GO:0004169:dolichyl−phosphate−mannose−protein manno...
GO:0010485:H4 histone acetyltransferase activity
GO:0001968:fibronectin binding
GO:0016634:oxidoreductase activity, acting on the C...
GO:0016812:hydrolase activity, acting on carbon−nit...
GO:0017046:peptide hormone binding
GO:0070061:fructose binding
Biological process
Molecular Function
0
1
2
3
4
−log10(p−value)
Downregulated 5−Aza genes
GO:0044281:small molecule metabolic process
GO:0017144:drug metabolic process
GO:0055114:oxidation−reduction process
GO:0019752:carboxylic acid metabolic process
GO:0006082:organic acid metabolic process
GO:0043436:oxoacid metabolic process
GO:0055086:nucleobase−containing small molecule met...
GO:0055085:transmembrane transport
GO:0072521:purine−containing compound metabolic pro...
GO:0006163:purine nucleotide metabolic process
GO:0016054:organic acid catabolic process
GO:0046395:carboxylic acid catabolic process
GO:0006753:nucleoside phosphate metabolic process
GO:0006839:mitochondrial transport
GO:0009117:nucleotide metabolic process
GO:0006790:sulfur compound metabolic process
GO:0009150:purine ribonucleotide metabolic process
GO:0034220:ion transmembrane transport
GO:0009063:cellular amino acid catabolic process
GO:0045333:cellular respiration
GO:0046034:ATP metabolic process
GO:0009259:ribonucleotide metabolic process
GO:1901617:organic hydroxy compound biosynthetic pr...
GO:0006119:oxidative phosphorylation
GO:0045471:response to ethanol
GO:0006091:generation of precursor metabolites and ...
GO:0009060:aerobic respiration
GO:0042026:protein refolding
GO:0006099:tricarboxylic acid cycle
GO:0044282:small molecule catabolic process
GO:0022857:transmembrane transporter activity
GO:0016491:oxidoreductase activity
GO:0022804:active transmembrane transporter activit...
GO:0005215:transporter activity
GO:0043177:organic acid binding
GO:0036094:small molecule binding
GO:0015075:ion transmembrane transporter activity
GO:0031406:carboxylic acid binding
GO:0003824:catalytic activity
GO:0043168:anion binding
GO:0043167:ion binding
GO:0016597:amino acid binding
GO:0000166:nucleotide binding
GO:1901265:nucleoside phosphate binding
GO:0031249:denatured protein binding
GO:0022853:active ion transmembrane transporter act...
GO:0020037:heme binding
GO:0015399:primary active transmembrane transporter...
GO:0004185:serine−type carboxypeptidase activity
GO:0051059:NF−kappaB binding
GO:0055131:C3HC4−type RING finger domain binding
GO:0015318:inorganic molecule entity transmembrane...
GO:0009055:electron transfer activity
GO:0070008:serine−type exopeptidase activity
GO:0005388:calcium transmembrane transporter activi...
GO:0046906:tetrapyrrole binding
GO:0097718:disordered domain specific binding
GO:0008236:serine−type peptidase activity
GO:0017171:serine hydrolase activity
GO:0005509:calcium ion binding
Biological process
Molecular Function
0
5
10
15
−log10(p−value)

**Fig. S8.**

**Transcriptional response to 5-Azacytidine in *Amoebidium* and *Sphaeroforma*.** (**A**) Upset plot showing the amount of differentially expressed (DE) genes (DEseq2, one to one orthologues) shared across *Amoebidium appalachense* and *Sphaeroforma arctica* comparing DMSO and 5-Azacytidine. (**B**) Transcriptional response of TE individual insertions to 5-Azacytidine treatment in *Amoebidium* and *Sphaeroforma* measured by TElocal. Only TEs larger than 500 bp were included. "No change" indicate TEs which do not change expression level (fdr > 0.05) or are silenced throughout. (**C**) Most abundant Pfam domains encoded in pandoravirus endogenized genes reactivated upon 5-Azacytidine treatment. (**D**) Gene ontology enrichments of *Amoebidium* host genes upregulated and downregulated upon 5-Azacytidine treatment.

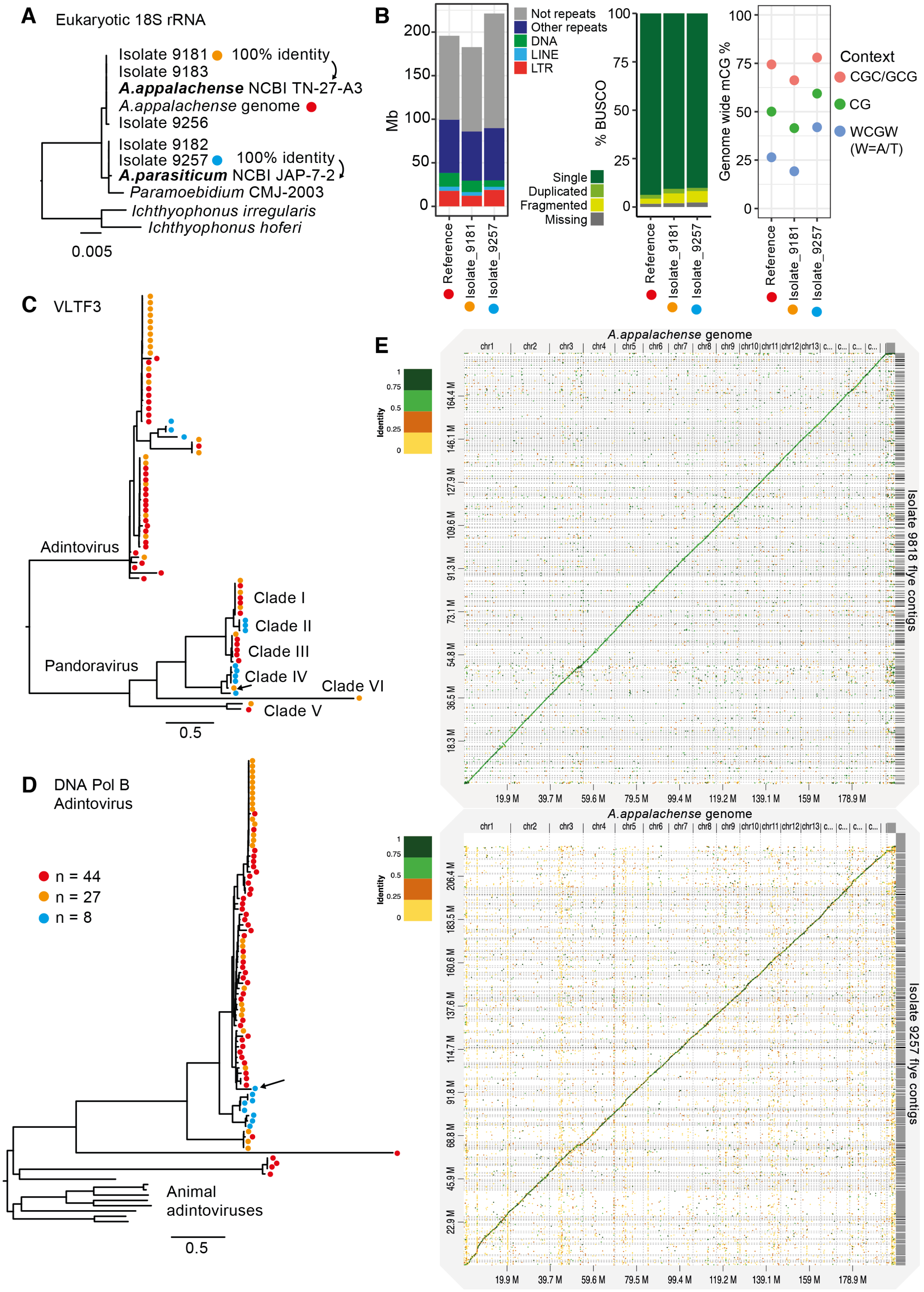
A
Eukaryotic 18S rRNA
Isolate 9181
100% identity
Isolate 9183
A.appalachense NCBI TN-27-A3
A.appalachense genome
Isolate 9256
Isolate 9182
Isolate 9257
100% identity
A.parasiticum NCBI JAP-7-2
Paramoebidium CMJ-2003
Ichthyophonus irregularis
Ichthyophonus hoferi
0.005
B
Mb
Not repeats
Other repeats
DNA
LINE
LTR
Reference
Isolate_9181
Isolate_9257
% BUSCO
Single
Duplicated
Fragmented
Missing
Genome wide mCG %
Context
CGC/GCG
CG
WCGW
(W=A/T)
C
VLTF3
Adintovirus
Pandoravirus
Clade I
Clade II
Clade III
Clade IV
Clade VI
Clade V
0.5
D
DNA Pol B
Adintovirus
n = 44
n = 27
n = 8
Animal
adintoviruses
0.5
E
A.appalachense genome
Isolate 9818 flye contigs
Isolate 9257 flye contigs
Identity

**Fig. S9.**

**Genomes and endogenized viral phylogenies of *Amoebidium* isolates.** (**A**) Maximum likelihood phylogenetic tree displaying the nuclear ribosomal 18S RNA phylogeny of *Amoebidium* isolates and closely related *Ichthyophonus* and *Paramoebidium* genus. The Isolate 9181 (orange) 18S is 100% identical to *A. appalachense* in NCBI and Isolate 9257 (blue) 18S is 100% identical to *A. parasiticum* in NCBI. (**B**) Genome size, repeat distribution, BUSCO completeness and global CG methylation levels across the 3 *Amoebidium* genomes. (**C**) Maximum likelihood phylogeny of VLTF3 encoded in pandoravirus and adintovirus endogenization events. Dots represent the genome they come from following panel **Fig. 4A** colour code. (**D**) Maximum likelihood phylogeny of DNA Polymerase family B encoded in adintoviruses. Coloured dots represent genome of origin. (**E**) Dot plot showing the genome alignment of contig level assemblies of Isolate 9181 and 9257 compared to the chromosome scale *A. appalachense* reference genome. Identity of alignment windows is calculated with minimap2 (with the D-GENIES server).
